# Long range mutual activation establishes Rho and Rac polarity during cell migration

**DOI:** 10.1101/2024.10.01.616161

**Authors:** Henry De Belly, Andreu Fernandez Gallen, Evelyn Strickland, Dorothy C. Estrada, Patrick J. Zager, Tamas L. Nagy, Janis K Burkhardt, Hervé Turlier, Orion D. Weiner

## Abstract

In migrating cells, the GTPase Rac organizes a protrusive front, whereas Rho organizes a contractile back. How these GTPases are appropriately positioned at the opposite poles of migrating cells is unknown. Here we leverage optogenetics, manipulation of cell mechanics, and mathematical modeling to reveal a surprising mechanochemical long-range mutual activation of the front and back polarity programs that complements their well-known local mutual inhibition. Rac-based protrusion stimulates Rho activation at the opposite side of the cell via membrane tension-based activation of mTORC2. Conversely, Rho-based contraction induces cortical-flow-based regulation of phosphoinositide signaling to trigger Rac activation at the opposite side of the cell. We develop a minimal unifying mechanochemical model of the cell to explain how this long-range facilitation complements local inhibition to enable robust Rho and Rac partitioning. We show that this long-range mutual activation of Rac and Rho is conserved in epithelial cells and is also essential for efficient polarity and migration of primary human T cells, indicating the generality of this circuit. Our findings demonstrate that the actin cortex and plasma membrane function as an integrated mechanochemical system for long-range partitioning of Rac and Rho during cell migration and likely other cellular contexts.

## Introduction

For proper physiology, many cells need to polarize, or restrict different signaling programs to different portions of their cell surface. Rac and Rho GTPases are key regulators of cell polarity and spatially pattern the actin cytoskeleton for mitosis, morphogenesis, migration, and development (1). Rac is localized to the front migrating cells, where it promotes the extension of sheet-like lamellipodia or pressure-driven blebs **(Figure 1A)** (2, 3). Rho is localized to the back of migrating cells, where it promotes local myosin-2-based contractions and long-range cortical flows (1, 4) **(Figure 1A)**.

**Fig. 1.**
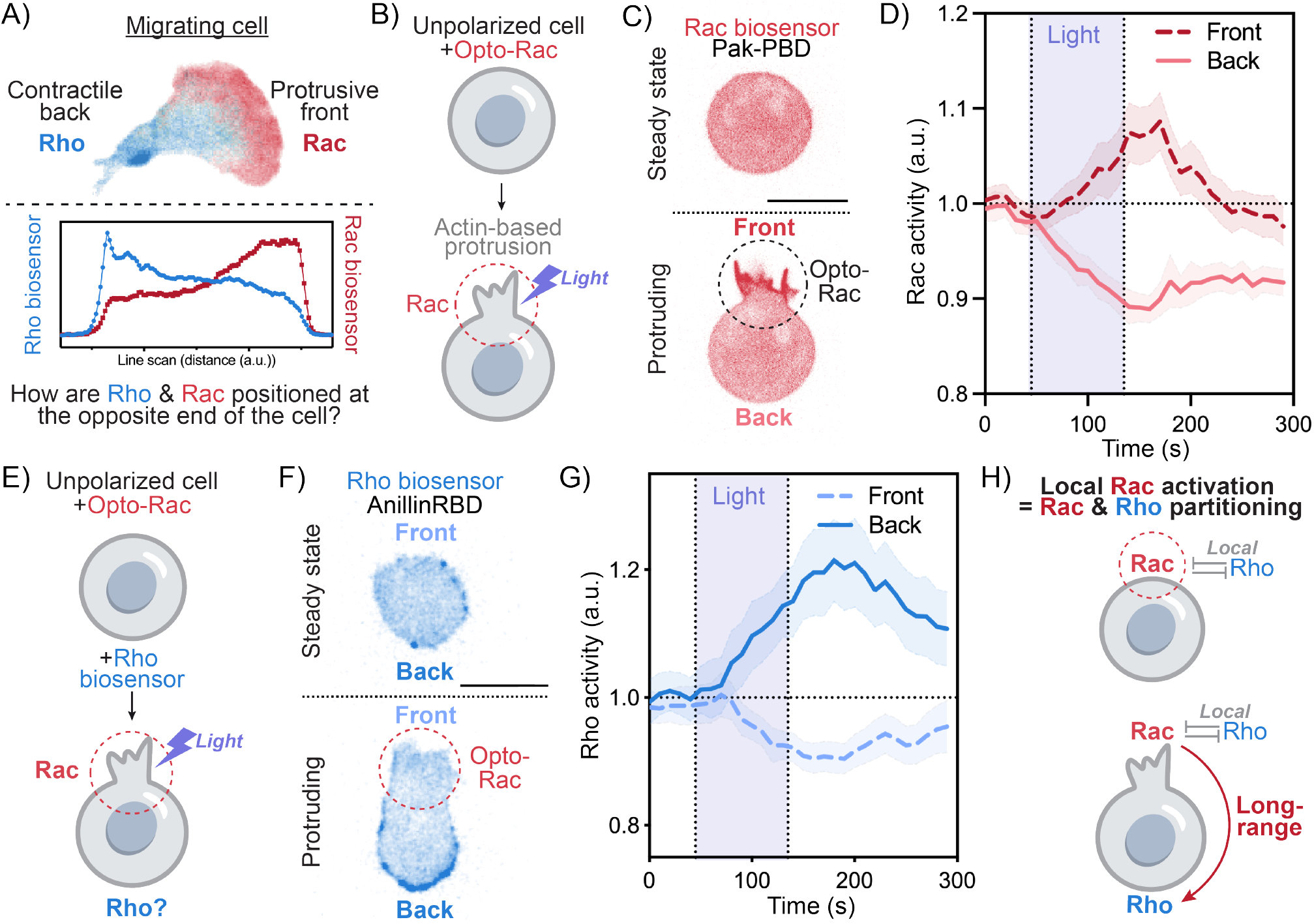
Local Rac activation stimulates long-range RhoA activation at the opposite side of the cell. (A) Confocal image and associated linescan of a migrating neutrophil-like HL-60 cell expressing the polarity biosensors for Rac (PAK-PBD, in red) and Rho (Anillin-RBD, in blue). In migrating cells, Rac and Rho localize to the protruding cell front and contracting cell back, respectively. Here we sought to investigate how Rac and Rho are appropriately positioned at the opposite poles of a migrating cell. (B) We locally activated the front polarity program Rac in an initially unpolarized neutrophil-like HL-60 cell using optogenetics (opto-PI3K, see Methods). (C) Time-lapse confocal images of an unpolarized cell before and during opto-Rac stimulation. Rac activity was monitored via the Rac biosensor Pak-PBD. (D) Average time trace of Rac activity at the plasma membrane at the site of opto-Rac activation (cell front) compared to the opposite side of the cell (cell back). (mean ± 95%CI; n>40, N= 3). (E) We locally activated Rac in an initially unpolarized cell while simultaneously measuring Rho activity using AnillinRBD. (F) Time-lapse confocal images of an unpolarized cell before and during opto-Rac stimulation. Rho activity was monitored using the biosensor AnillinRBD. (G) Average time trace of Rho activity at the plasma membrane for the front versus back of the cell following opto-Rac activation. (mean ± 95%CI; n>40, N= 8). (H) Local Rac activation leads to long-range activation of Rho at the opposite side of the cell. Scale bars: 10*µ*m.

While we know that Rho and Rac mutually inhibit each other locally (1, 5–7), how they are properly positioned at opposite ends of the cell during migration remains poorly understood. The majority of models of polarity in migration are based on local biochemical interactions (6, 8–15), with a majority of mathematical models of polarity establishment relying in fact on the so-called wave-pinning mechanism (16–19), where a travelling front of active Rho GTPase is pinned by a bath of cytosolic inactive form. Such mechanism relies importantly on the fast diffusion of the inactive form and a hypothesis of fixed total protein amount. However, previous studies have challenged these requirements, demonstrating that a diffusion-based inhibitor or depletion mechanism is not sufficient to explain neutrophil polarization (20–23). In contrast, local inhibition is not sufficient either to explain the long-rang partitioning of active pools of Rho and Rac and to coordinate front and back at the scale of a migrating cell. The requirement for an additional long-range communication had long been suggested (20–25), but direct evidence and molecular mechanism for this link was unknown.

Cell polarity involves long-range cellular information processing and necessitates information flow at the cellular scale. Forces transmitted via the actin cortex and the plasma membrane have emerged as key conduits for this global coordination. In migration, membrane tension guides shape by relaying actin-based protrusive forces at the front to the contraction of the rear; setting up a global competition that enables the establishment of a single dominant front (26–30). Here we investigate whether the membrane and cortex could act as the mechanical conduit for long-range coordination of the front and back polarity programs.

To probe the long-range spatial coordination of Rac and Rho, we take advantage of optogenetic activators of Rac and Rho in initially unpolarized cells. This enables us to precisely and specifically activate either Rac or Rho in a small region of the cell and observe the global response of the other GTPase. By combining optogenetics with manipulation of cell mechanics and mathematical modeling, we find that both the front and back mutually activate each other at a distance, and this occurs using two distinct pathways. The front stimulates the back via membrane tension, whereas the back stimulates the front via cortical remodeling. We demonstrate the physiological relevance of our findings for immune cell migration using primary human T cells. Our results demonstrate that the actin cortex and plasma membrane act as an integrated mechanochemical system to ensure the proper positioning of the front and back polarity programs during cell migration.

## Results

### Local Rac activation stimulates long-range RhoA activation at the opposite side of the cell

To investigate how the front and back polarity programs are properly positioned at the opposite ends of the cell, we first analyzed the establishment of cell polarity in a context where we can independently control the spatial and temporal dynamics of each of these programs. Towards this end, we leveraged an optogenetic approach (via local activation of PI3K (31), see Methods), to locally activate the front polarity program Rac in initially unpolarized neutrophil-like HL60 cells **(Figure S1A)**. As previously demonstrated, this local Rac activation leads to actin-driven protrusions like those seen at the cell front during migration **(Figure S1)** (31, 32). Rac activity (visualized with Pak-PBD) is locally enriched in the zone of activation (cell front), while being depleted at the opposite side of the cell (cell back) **(Figure 1B-D, S1B, C and Video S1)**. These results confirm that opto-PI3K locally activates Rac. Next, we sought to investigate how this local zone of Rac activity influences the back polarity program. To this end, we used local Rac activation via opto-PI3K while monitoring the activity of Rho using the biosensor Anillin-RBD **(Figure 1E and Video S2)**. Light-induced Rac activation elicits a rapid long-range increase in Rho activity at the opposite side of the cell **(Figures 1F,G and S1G-I)**. Thus, in addition to locally inhibiting Rho, Rac activation also stimulates Rho at the opposite end of the cell.

We next sought to understand how local Rac activation in one part of the cell triggers Rho activity at the opposite end. We considered two possibilities: distal Rho activation could result from local Rac-based biochemical inhibition(1) or indirect long range mechanical signals. One possible means of long-range Rho activation would be Rac-induced protrusions, which generate a membrane tension increase that propagates across the cell (32) **(Figure S2A)**. To distinguish between these possibilities, we impaired Rac-mediated protrusion formation by applying pharmacological inhibitors of actin assembly (latrunculin) or Arp2/3 complex activation (CK666) to opto-PI3K cells **(Figure S2A-D)**. While these perturbations do not impair our ability to optogenetically activate Rac (31), they inhibited Rac-mediated Rho activation at the other end of the cell. To test whether Rho activation depend on the actomyosin network, we impaired actomyosin contractility by treating cells with either the myosin inhibitor blebbistatin or the ROCK inhibitor Y-27632 **(Figure S2E)** and observed no noticeable effect on Rho as a result of local Rac activation **(Figure S2F-H)**. These data indicate that protrusion mediated Rho activation in independent of contractility. Altogether, our results support long-range activation of Rho by Rac, operating independently from the well-established local antagonism between these GTPases **(Figure 1H)**.

### Rac stimulates long-range Rho activation via membrane-tension-mediated mTORC2 activation

If protrusions stimulate Rho activation through an increase in membrane tension (32) **(Figure 2A)**, membrane tension increases should suffice to stimulate Rho activation even in the absence of Rac activation or protrusions. To investigate this possibility, we used hypotonic shock to stimulate an increase in membrane tension (33) **(Figure 2B)**. Hypotonic shock induced a rapid and global increase in Rho activity across the cell **(Figures 2C, D, S3A-F and Video S3)**. To verify that this increase in Rho activity is a function of increased membrane tension and not a secondary consequence of the rebuilding of the actin cortex following hypotonic shock, we combined our hypotonic shock assay with Latruculin B treatment, a combination that potently depolymerizes the actin cortex (32, 33). Under these conditions, osmotic shock still activates Rho, indicating that the actin cytoskeleton is not necessary for membrane-stretch-based Rho activation **(Figure S3D)**. As an alternate approach, we next used a micropipette-based aspiration assay, which we previously demonstrated to increased membrane tension32. This mechanical manipulation also simulates global Rho activation **(Figures 2E-G; S3G-J and Video S4)**. The activation of Rho by hypotonic shock, micropipette aspiration and opto-induced protrusions likely reflects a role for membrane tension as a Rho activator. To test whether increases in tension also affect Rac activation, we performed measurements of Rac activity in response to both osmotic shock and micropipette aspiration and found that, as previously reported, elevated tension does not activate Rac **(Figure S3E, F and I, J)**. Our data show that Rac acts through protrusion-mediated increases in membrane tension to stimulate Rho at a distance.

**Fig. 2.**
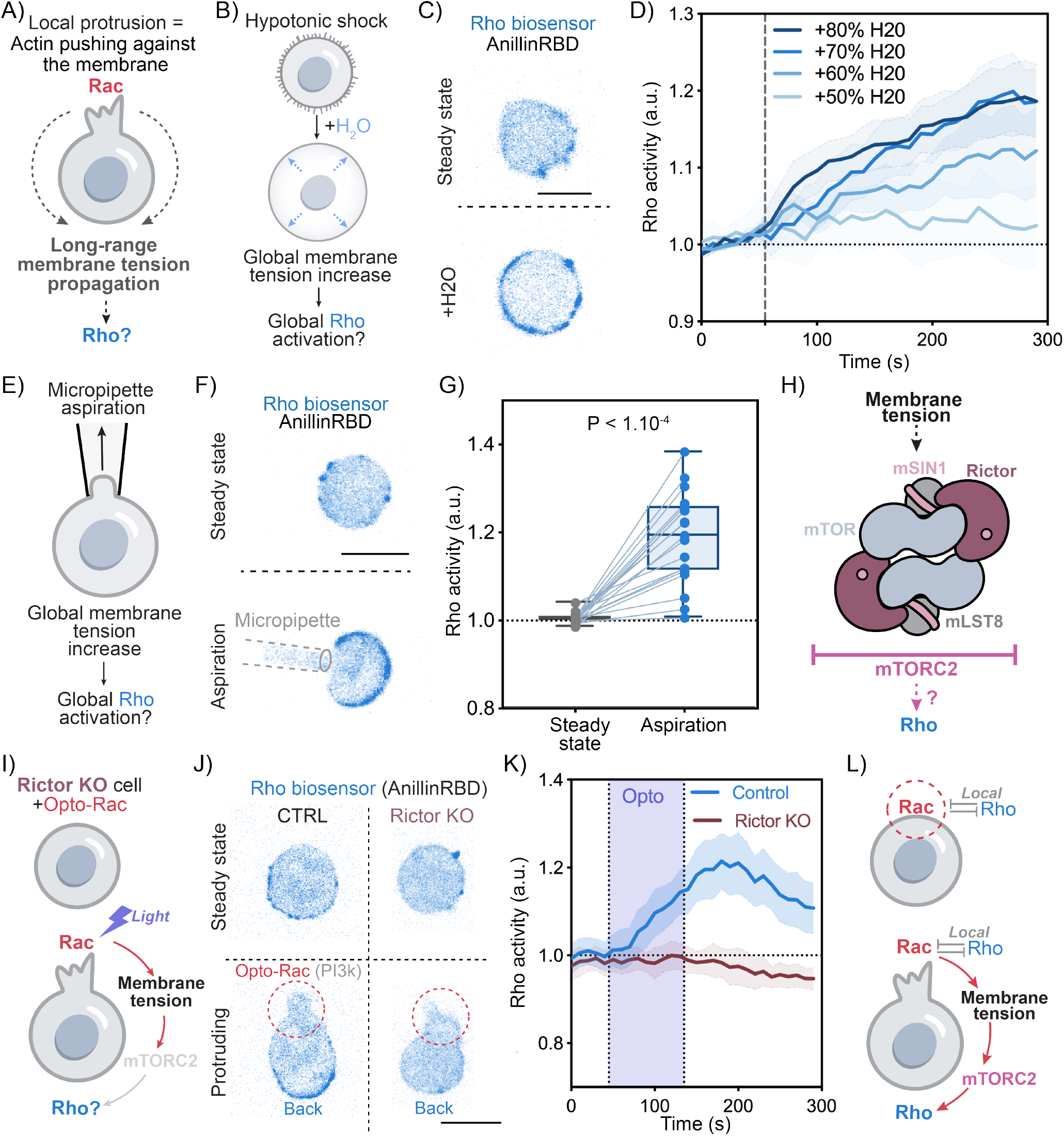
Rac stimulates long-range RhoA activation at the opposite side of the cell via membrane-tension-mediated mTORC2 activation. (A) Local Rac-mediated cell protrusion leads to a global increase in membrane tension32. (B) If protrusions activate Rho by elevating membrane tension, then increasing membrane tension (even in the absence of Rac activation) should suffice to activate Rho. Towards this end, we used hypotonic shock to elevate membrane tension. (C) Time-lapse confocal images of an unpolarized opto-Rac HL-60 cell expressing the Rho biosensor (AnillinRBD) before and after hypotonic shock. (D) Average time trace of Rho activity at the plasma membrane in response for hypotonic shocks of various intensity. (N>3, n>20, means ± 95%CI). Hypotonic-shock-based elevation of membrane tension suffices to globally increase RhoA activity. (E) As an alternate approach to increase membrane tension, we leveraged micropipette aspiration(32). (F) Time-lapse confocal images of an unpolarized opto-Rac HL-60 cell expressing the Rho biosensor (AnillinRBD) before and after micropipette aspiration (see Methods). (G) Average Rho activity before (steady state) and during aspiration. Micropipette-based elevation of membrane tension significantly stimulates Rho activation. Box and whiskers: median and min to max; p values from Wilcoxon paired Student’s t test. (H) We investigate whether mTORC2 is part of the mechanosensory pathway that links increased in membrane tension to the activation of Rho. (I) We used optogenetics to locally activate Rac in control cells versus cells lacking different core components of the mTORC2 complex (Rictor or mSIN1 knockouts). (J) Time-lapse confocal images of an unpolarized control or Rictor KO cell before and during opto-Rac stimulation. Rac activity was monitored via the Rac biosensor Pak-PBD. (K) Local Rac activation potently stimulates long-range activation of Rho in control cells (blue) but not cells deficient in mTORC2 activity (Rictor KO, magenta) (mean ± 95%CI; n>20, N= 3). Control curve is same as in Fig. 1D. (L) Local Rac activation stimulates long-range Rho activation via membrane tension mediated mTORC2 activation. Scale bars: 10*µ*m.

Next, we sought to identify the mechanosensors that link increases in membrane tension to Rho activation. The mTORC2 complex emerged as a strong candidate, given its responsiveness to membrane tension and its key roles in regulating cell polarity and motility (28, 29, 33–35)**(Figure 2H)**. To test whether mTORC2 links membrane tension increases to Rho activation, we performed our opto-Rac activation assay in cells lacking different core components of the mTORC2 complex (Rictor or mSIN1 CRISPR KO cell lines) (29) or using a pharmacological inhibition of mTOR. Cells deficient in mTORC2 (Rictor or mSIN1 KO) or treated with mTOR inhibitor KU-0063794 failed to activate Rho activity upon local Rac-activation **(Figures 2I-K; S4A-E and Video S5)** despite retaining their ability to generate protrusions. To further isolate the tension-mediated potentiation of Rho activity, we repeated previous hypotonic and micropipette aspiration assays in mTORC2-impaired cells. Our results confirmed that mTORC2 is essential for Rho activation following both hypotonic shock **(Figures S3F-H)** and micropipette aspiration **(Figures S3I-K)**. Our work reveals a molecular pathway in which Rac-induced actin protrusions globally increase membrane tension, activating the mechanosensitive mTORC2, which in turn stimulates Rho.

### Local Rho activation leads to Rac activation at the opposite side of the cell

We previously established that local Rac-mediated protrusion is sufficient for long-range Rac and Rho partitioning across the cell. However, not all migrating cells generate branched actin-based protrusions at their front. In bleb-based migration, Rho mediated contraction at the cell back is the primary force generator, producing increased intracellular pressure leading to blebbing on the opposite side (2, 36). Interestingly, while both blebbing programs and actin polymerization-based protrusions use different protrusion engines, both rely on the same Rho-Rac polarity machinery (37, 38). Given the role of Rho activation in promoting bleb-based migration, we next wondered whether local Rho activation is also sufficient to trigger long-range Rac and Rho partitioning.

To investigate the ability of Rho to activate Rac, we used an optogenetic approach to locally activate the back polarity program Rho (via LARG, see Methods) while simultaneously monitoring Rac activity using the biosensor Pak-PBD **(Figure 3A and Video S6)**. Our results show that Rho activation leads to long-range Rac activation at the opposite end of the cell, coinciding with another morphological change— blebbing **(Figure 3B)**. Given previous studies linking blebbing to Rac activation (38, 39), we hypothesized that Rho’s long-range stimulation of Rac might depend on Rho-induced blebbing **(Figures 3C-E and S5A)**. To test this hypothesis, we used the actomyosin inhibitor blebbistatin to block cellular contraction and blebbing following optogenetic Rho activation **(Figure 3F)**. Indeed, when blebbing was inhibited, Rho could no longer activate Rac at a distance **(Figures 3G, H)**. Just as Rac requires its downstream cytoskeletal effects (actin polymerization and protrusion generation) to activate Rho at a distance **(Figure S2)**, Rho relies on its downstream cytoskeletal impact (myosin-based contraction and blebbing) to activate Rac at a distance. To further explore whether blebs suffice to activate Rac independent of Rho-mediated stimulation of actomyosin contractility, we used the actin inhibitor Latrunculin B to induce stable blebs (32) **(Figure S5B)**. These stable blebs showed robust Rac enrichment **(Figure S5C, D)**. Our data demonstrate that local Rho activation stimulates Rac activity at the opposite end of the cell through actomyosin-based blebbing.

**Fig. 3.**
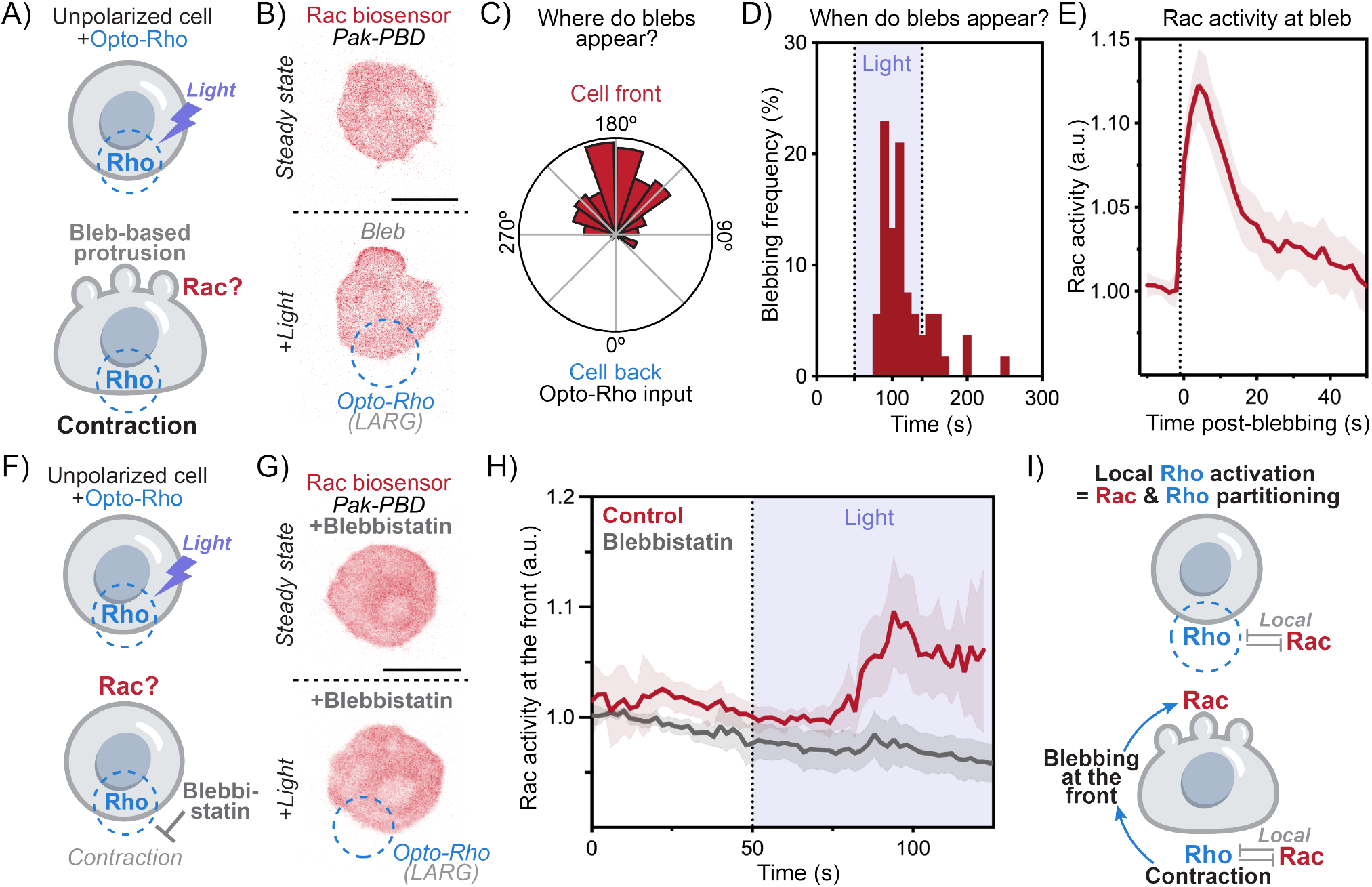
Local Rho activation elicits Rac activation at the opposite side of the cell. (A) Local Rac activation leads to long-range Rho activation at the opposite end of the cell via membrane tension. Does local Rho activation also lead to long-range Rho and Rac partitioning? To test this, we used optogenetics to locally stimulate Rho (opto-LARG, see Methods) while simultaneously measuring Rac activity using the biosensor Pak-PBD. (B) Time-lapse confocal images of an unpolarized opto-Rac HL-60 cell expressing the Rac biosensor (Pak-PBD) before and during opto-Rho stimulation. Local Rho activation induces blebs at the opposite end of the cell, and these blebs are enriched in active Rac. (C) Polar histogram of the spatial distribution of blebs appearance in relation to opto-Rho (cell back) showing that the majority of blebs are positioned at the opposite side of the cell back (opto-Rho). (N = 5, n > 65). (D) Histogram of bleb distribution in time during opto-Rho stimulation. (N = 3, n>55). (E) Average time trace of Rac activity at the plasma membrane at the protrusive blebs. Time 0 = blebbing. For times before blebbing, Rac activity is measured at the same membrane spot where the bleb will appear. (see Methods) (mean 95%CI; n > 30, N = 3). (F) We hypothesized that Rho’s long-range stimulation of Rac might depend on Rho-based contractility which in turns triggers blebbing. (G) We locally activated Rho using optogenetics in the absence or presence of an inhibitor of myosin activation and blebbing (10*µ*M Blebbistatin). (H) Average time trace of Rac activity at the plasma membrane measured at the cell front of control or cells treated with 10M Blebbistatin. These data indicate that myosin-based blebbing is important for Rho-mediated activation of Rac activity (mean ± 95%CI; n>20, N= 3). (I) Local Rho activation leads to long-range activation of Rac at the opposite side of the cell. Scale bars: 10*µ*m.

### Contraction-induced membrane-to-cortex attachment asymmetry leads to PIP2 release and PI3K-dependent Rac activation at the opposite side of the cell

We found that Rac is recruited to blebs opposite the contractile back of the cell, but what determines the position of these blebs relative to the Rho-mediated contraction site? Blebs form when the plasma membrane detaches from the underlying actin cortex, and this process is facilitated by a low local concentration of membrane-to-cortex attachment (MCA) proteins. Localized contraction induces actomyosin flow toward the site of Rho mediated-contraction (32). This flow could cause MCA proteins to accumulate at the site of contraction while depleting them from the opposite end of the cell(40–42) **(Figure 4A)**. To test this hypothesis, we imaged cells expressing a fluorescent tagged ezrin, a core MCA protein in neutrophils. Upon local Rho activation, ezrin accumulated at the site of contraction and was depleted at the opposite side of the cell **(Figure 4B, C; S6A-C and Video S7)**. To further validate these findings, we used actin-membrane proximity biosensor MPAct (43) to assess MCA distribution during opto-contraction. This tool confirmed that local contraction enriches MCA at the contraction site while depleting it from the opposite side of the cell **(Figure S6D-F)**. These results suggest that the depletion of MCA opposite to the contraction site promotes the formation of bleb-based sites of Rac activation.

**Fig. 4.**
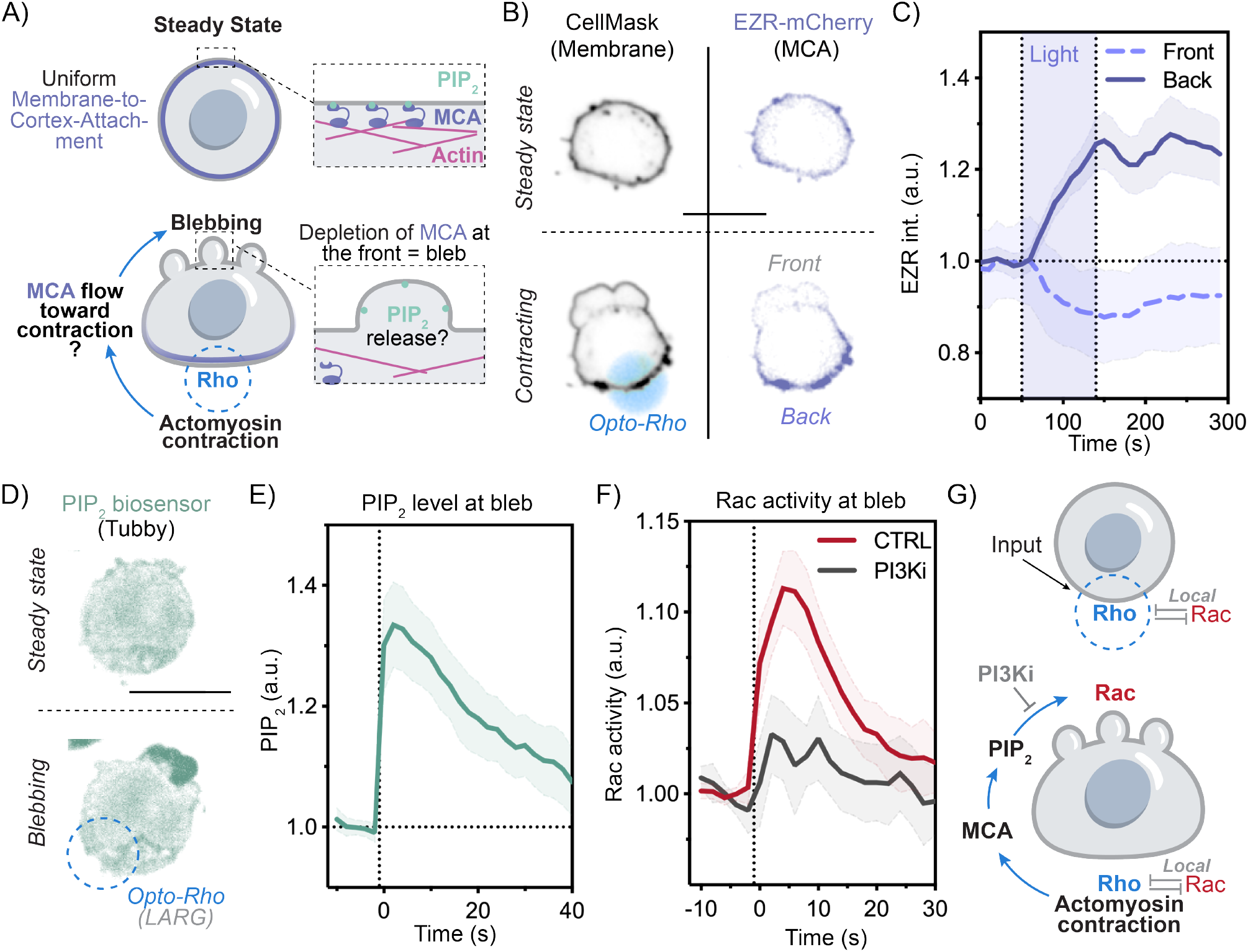
Contraction-induced blebbing leads to PIP2 release and PI3K-dependent Rac activation at the opposite side of the cell. (A)In unpolarized cells at steady state, actin and MCA proteins are uniformly distributed. Upon local contraction (for example induced by opto-Rho), Actin flows toward the site of contraction. We ask whether MCA also flow with the cytoskeleton toward the site of contraction to induce an MCA asymmetry across the cell. (B) Time-lapse confocal images of unpolarized cells expressing the MCA protein Ezrin-mCherry and stained with the membrane marker CellMask before and during opto-Rho stimulation. (C) Average time trace of relative Ezrin intensity at the back (purple) and front (grey) of the cell in response to opto-Rho activation (at the back). (N = 3, n >40, means ± 95%CI). We next hypothesized that MCA detachment from plasma membrane during blebbing leads to PIP2 release. (D) Time-lapse confocal images of an unpolarized opto-Rac HL-60 cell expressing the PIP2 biosensor (Tubby) before and during opto-Rho stimulation demonstrates PIP2 enrichment in bleb-based protrusions. (G) Average time trace of PIP2 accumulation at the protrusive blebs (opposite side of the cell from Rho activation) (mean ± 95%CI; n>30, N= 3). (I) Average time trace of Rac activity at the plasma membrane at the protrusive blebs in control and cells treated with 1 *µ*M PI3Kδ/γ inhibitor, Duvelisib demonstrates a requirement for PI3K for Rac activation in protrusive blebs (mean ± 95%CI; n>30, N= 3). Control data is the same as in Figure 3E. (G) Local Rho activation leads to contraction-driven localized PIP2 enrichment at the cell front which act a substrate of the Rac activator PI3K. Scale bars: 10*µ*m.

We next explored the molecular mechanism linking blebbing to Rac activation, hypothesizing the lipid composition of the blebs to play a critical role. Because MCA proteins such as ERMs (Ezrin, Radixin, Moesin) bind to the membrane through interactions with PIP2 (44), detachment of MCA proteins from the membrane during blebbing could increase the local concentration of accessible PIP2. PI3K could act on this newly-released PIP2 to produce PIP3, a known activator of Rac. To test this hypothesis, we used the PIP2 biosensor Tubby (45) in conjunction with local opto-Rho activation and observed a marked increase in PIP2 level at blebs induced by Rho mediated contraction **(Figure 4D, E and Video S8)**. Next, we examined the functional role of this PIP3 in Rac activation within blebs by inhibiting PI3K using Duvelisib (PI3Ki), a dual PI3Kδ/γ inhibitor (46) **(Figures 4F and S6G, H)**. Cells treated with Duvelisib displayed a significant impairment Rac enrichment downstream of both opto-Rho-mediated blebs **(Figure 4F)** as well as Latrunculin-induced stable blebs **(Figure S6I**,**J)**. These findings provide a molecular mechanism to link local Rho activation to Rac activation at the opposite end of the cell. Local Rho-induced contraction redistributes MCA proteins away from the opposite end of the cell, promoting blebbing. This depletion of MCA proteins releases PIP2, which serves as a substrate for PI3K to generate the Rac activator PIP3**(Figure 4G)**.

### Core components of the mutual activation Rho and Rac activation are conserved in non-migratory cells

Our experiments demonstrate that Rho and Rac mutually activate one another at a distance using two distinct mechanisms: (1) membrane tension, which links front generation with back activation, and (2) cortical remodeling, which connects back activation with front generation. Given the broad conservation of the components of the pathways we have identified for this long-range coupling (membrane tension propagation, mTORC2 activation, cortical flows, and phospholipid changes), we next investigated the generality of this circuit beyond cell migration. First, we probed whether tension/mTORC2-mediated increases in Rho activity via mTORC2 is conserved in non-migratory cells **(Figure 5A)**. To investigate this possibility, we used hypotonic shock in resuspended epithelial cells (Hek-293Ts and HeLa) to stimulate an increase in membrane tension **(Figure 5B)**. Consistent with our previous findings, we observed that hypotonic shock induced a rapid and global increase in Rho activity across the cell **(Figures 5C, D)**. Cells treated with mTOR inhibitor KU-0063794 failed to activate Rho activity in response to hypotonic shock. These data show a conservation of the pathway linking membrane tension to mTORC2 to Rho activation in non-migratory cells **(Figures 5D and S7)**.

**Fig. 5.**
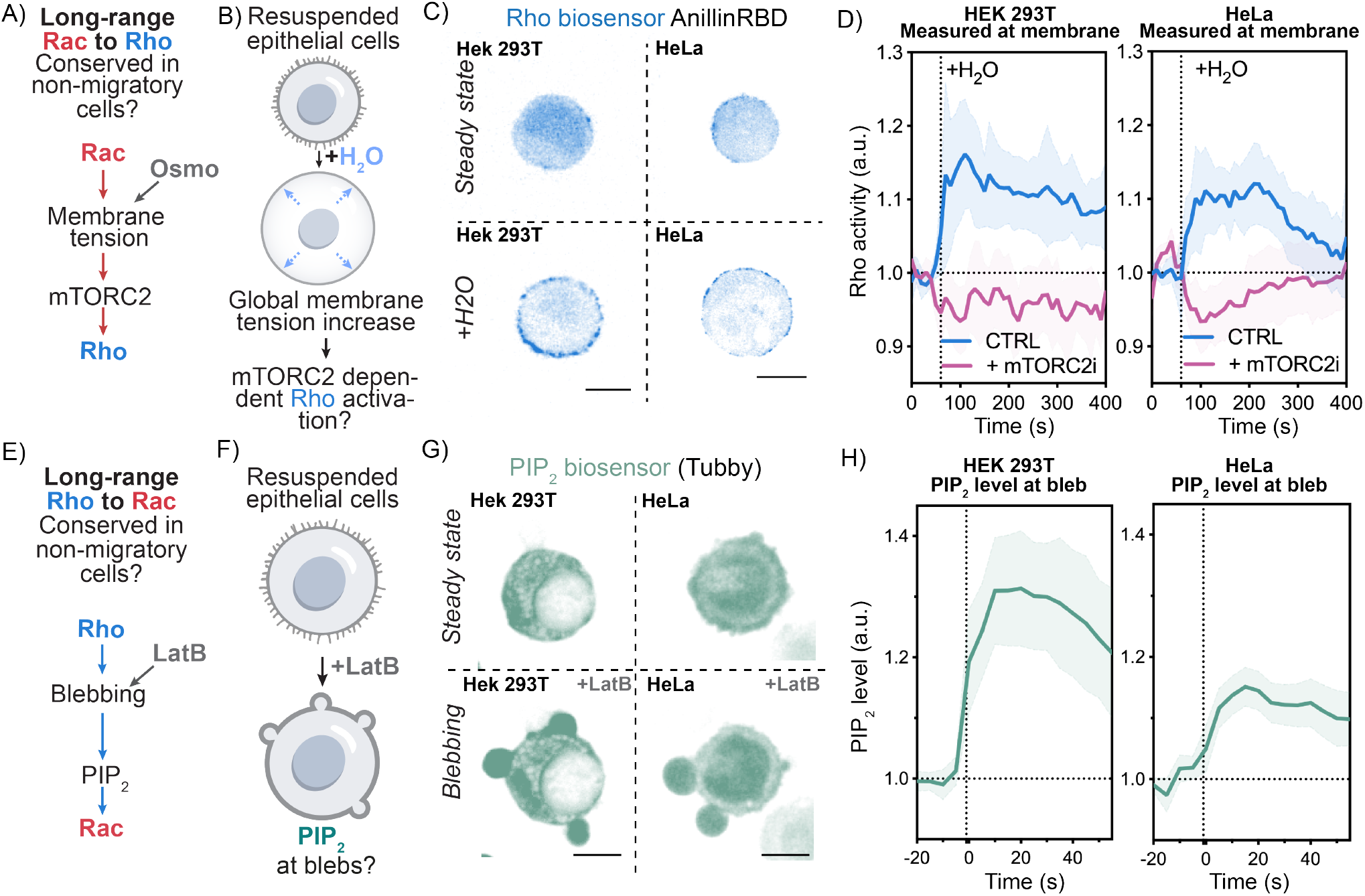
Core components of Rho and Rac mutual long-range activation are conserved in non-migratory cells. (A)Are the core components of the long-range communication from Rac to Rho conserved in non-migratory cells? (B) If membrane tension operates through mTORC2 to activate Rho in epithelial cells, then increasing membrane tension should suffice to activate Rho. Towards this end, we used hypotonic shock to elevate membrane tension in resuspended epithelial cells. (C) Confocal images of HeLa and 293Ts cells expressing the Rho biosensor Anillin-RBD before and during osmotic shock. (D) Average time trace of Rho activity at the plasma membrane in response for hypotonic shocks of control cells (in blue) or cells treated with the 10*µ*M of the mTOR inhibitor KU-0063794 (in pink) (N>2, n>20, means ± 95%CI). (E) Are the core components of the long-range communication from Rho to Rac conserved in non-migratory cells? (F) To test whether localized PIP2 release in blebs is conserved in non-migratory cells, we used resuspended epithelial cells treated with the actin inhibitor Latrunculin B to generate persistent cellular blebs. (G) Time-lapse confocal images of an unpolarized Hela and 293T cells expressing the PIP2 biosensor (Tubby) before and during Latrunculin B treatment demonstrates that PIP2 enrichment in bleb-based protrusions is conserved in non-migratory cells. (H) Average time trace of PIP2 accumulation at the blebs in HeLa and 293T cells treated with 2M of the actin inhibitor Latrunculin B (mean ± 95%CI; n>30, N= 2).

Next, we probed whether the MCA mediated release of PIP2 in blebs is also conserved in non-migratory cells **(Figure 5E)**. Towards this end, we used the actin inhibitor Latrunculin B to induce stable blebs in resuspended epithelial cells expressing the PIP2 biosensor Tubby **(Figure 5F)** and observed a marked increase in PIP2 in blebs in both HeLa and Hek 293T cells **(Figure 5G, H)**. Together, our results show that the core components of the long-range mutual activation between Rac and Rho are conserved in non-migratory cells and are likely to account for the robustness of polarity in a number of other cellular contexts, such as embryo polarization, epithelial polarity, and asymmetric cell division, all of which involve long-range Rac/Rho patterning.

### Mechanochemical model of Rho and Rac partitioning in cells

To understand the key requirements for the long-range Rho and Rac mutual activation, we developed a minimal mechanochemical model of Rho and Rac partitioning. We first formulated a mathematical model using two coupled differential equations to describe the local mutual inhibition between Rac and Rho (47) **(Figure 6A, Supplementary Text)**. The model defines two regions of stability for Rac and Rho, depending on their respective levels of antagonism-modulated activation, denoted *α*0 and ß0. At low or highly asymmetric production levels, only Rho or Rac will dominate (monostable regime), while at sufficiently high levels of both GTPases, a bistable regime arises where they can coexist **(Figure 6B, C)**. In the bistable regime, one could force the system locally to be either in a high Rac or Rho state, but polarization cannot persists in this simplified biochemical picture only **(Supplementary Text, section 2.4)**. This highlights the limitations of polarity regulation based only on local inhibition, consistent with our previous experimental results **(Figure S2A-D and 3F-H)**.

**Fig. 6.**
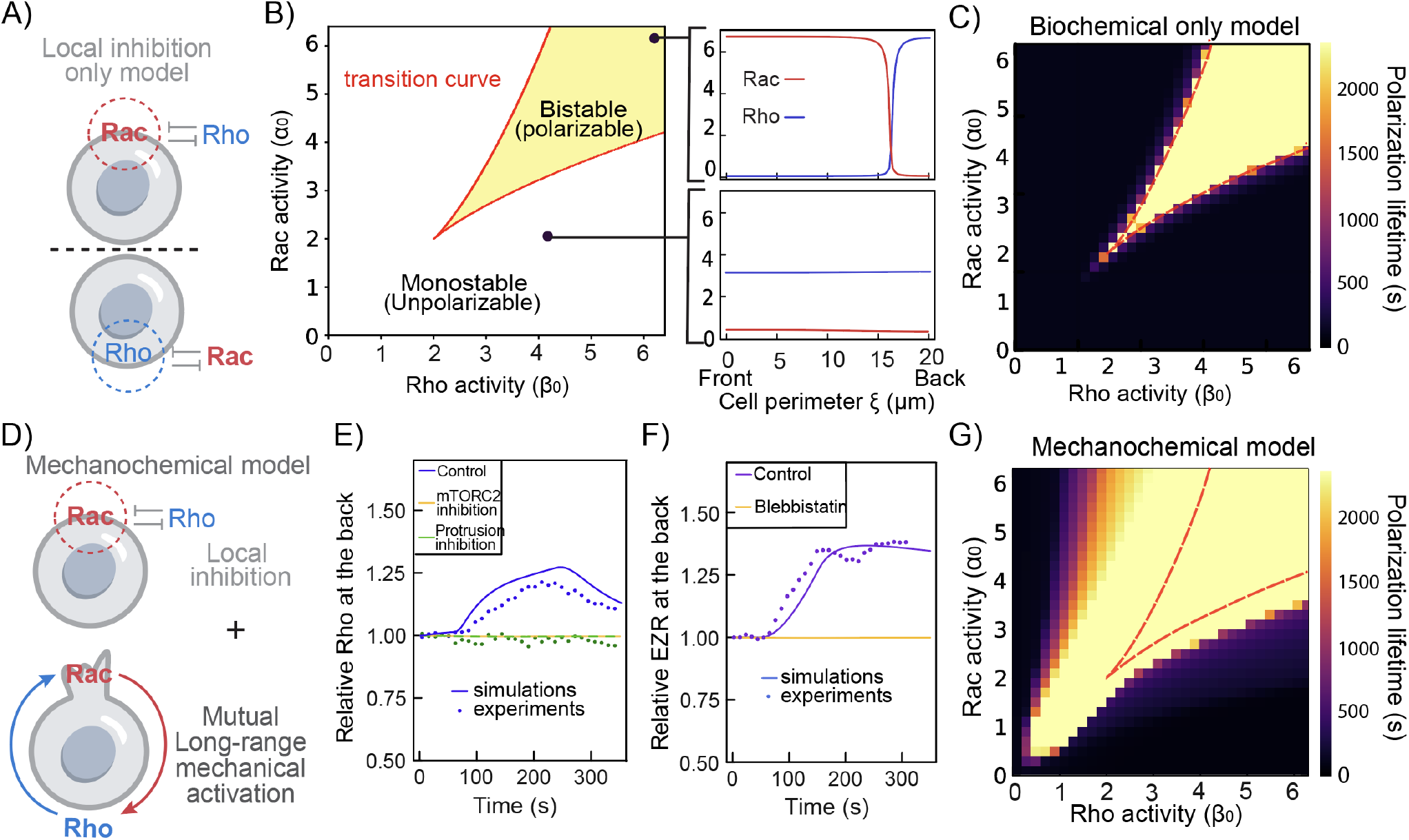
Modelling Rac and Rho polarization mechanisms through local inhibition and long-range mechanical feedback. (A)Simple model of local inhibition between Rac and Rho (B) Phase diagram of polarity obtained from stability analysis using a simple model of local mutual inhibition between Rac and Rho. For any given Rac and Rho basal levels (*α* and ß respectively), the system is either monostable (unpolarized) or bistable (polarized). On the right, concentration values of Rac and Rho for simulations inside (Top) and outside (Bottom) the bistable region, where (Top) is polarized with a high value of Rac at the front of the cell and low at the back and vice-versa for Rho. (C) Heatmap of the time needed for a polarized cell to relax back to steady state in function of basal Rac and Rho activity (*α* and ß respectively) using local inhibition only. (D) Mechanochemical model of Rac and Rho polarity, combining local mutual inhibition with long-range mechanical feedback. (E) Rac is locally activated on one end (the ‘front’) of the cell, while Rho activity is measured at the opposite end (the ‘back’) of the simulated cell. Simulated (solid lines) and experimental (dots) Rho activity at the back of the cell following Rac activation at the front. Control cells are shown in blue, mTORC2 defective cells (Rictor KO) in orange, cells with inhibited protrusions (CK666) in green. (F) Simulated (solid lines) and experimental (dots) Ezrin levels at the back of the cell show that local Rho activity leads to MCA asymmetry. Control is shown in purple while cells with impaired contraction (blebbistatin) are shown in yellow. (G) Same as (C) but using fully integrated mechanochemical model.

Next, we combined this mutual local inhibition model with a mechanical model describing the membrane-cortex interplay, building on previous works (32) **(see Supplementary Theory, section 4) (Figure 6D)**. In this model, the cortex is treated as a viscous-contractile layer attached to the membrane via MCA proteins of varying surface density ρ. The mechanically relevant membrane tension (also referred to as frame tension (48, 49)) is stored in entropic and cortical-flow-driven membrane ruffles, which are minimally modeled as a linearly elastic component. Rac is assumed to promote protrusion, modeled by a velocity (vp) at the front, while Rho mediates contraction at the back (σa). Based on previous experimental findings, we further introduce two novel couplings: activation of active Rac is negatively modulated by MCA protein density, and Rho is enhanced by membrane tension, each governed by switch-like functions. Using this mechanochemical model, we successfully replicated our previous findings showing where local opto-Rac activation resulted in Rho activation at the cell’s opposite **(Figure 6E and S7)**. In line with our experiments, blocking Rac’s influence on membrane tension (e.g., via drugs inhibiting protrusion) or disrupting tension’s effects on Rho activation (e.g., in mTORC2 mutants) prevented Rac from stimulating Rho activation at the distal end. This integrated mechanochemical model explains how the front positions the back by combining local inhibition and long-range, tension-mediated Rho activation. We next simulated the reverse scenario by locally activating Rho and measuring Rac act the opposite end. The model captured how Rho-driven creates MCA asymmetry **(Figure 6F)** and induces Rac activation at the distal pole **(Figure S7)**. The minor discrepancy between model and data for Rac activation may be due to variability in blebbing across cells, both in timing **(Figure 3D)** and quantity **(Figure S5A)**.

Finally, we compared the persistence of polarity generated by the integrated mechanochemical model with that produced by two alternative models: one based solely on local Rac-Rho mutual inhibition, and another using wave-pinning dynamics (16–19) driven by diffusion and mass conservation **(Supplementary Theory, Section 3)**. Our results indicate that local mutual inhibition alone is insufficient to robustly partition Rho and Rac across the cell **(Supplementary Theory, Figure 5)**. Although wave-pinning models can produce stable polarization, they exhibit hypersensitivity and a binary response to external stimuli or fluctuations in Rho and Rac—features that are inconsistent with the broader spectrum of cellular behaviors observed experimentally. In contrast, the integrated model, which combines local mutual inhibition with long-range mutual activation via mechanochemical coupling, successfully recapitulates experimentally observed behaviors. Specifically, it predicts three distinct regimes of cellular behavior across a wide range of Rac and Rho basal production levels: (1) a responsive but non-polarizable state, characteristic of HL60 cells; (2) a responsive and polarizable state, in which sufficiently strong stimulation can induce persistent polarization through mechanochemical feedback, consistent with the response of primary human T cells **(Figure 7)**; (3) a spontaneously polarized state, where the cell maintains a persistent polarized configuration without external cues **(Supplementary Theory, Figure 10)**.

**Fig. 7.**
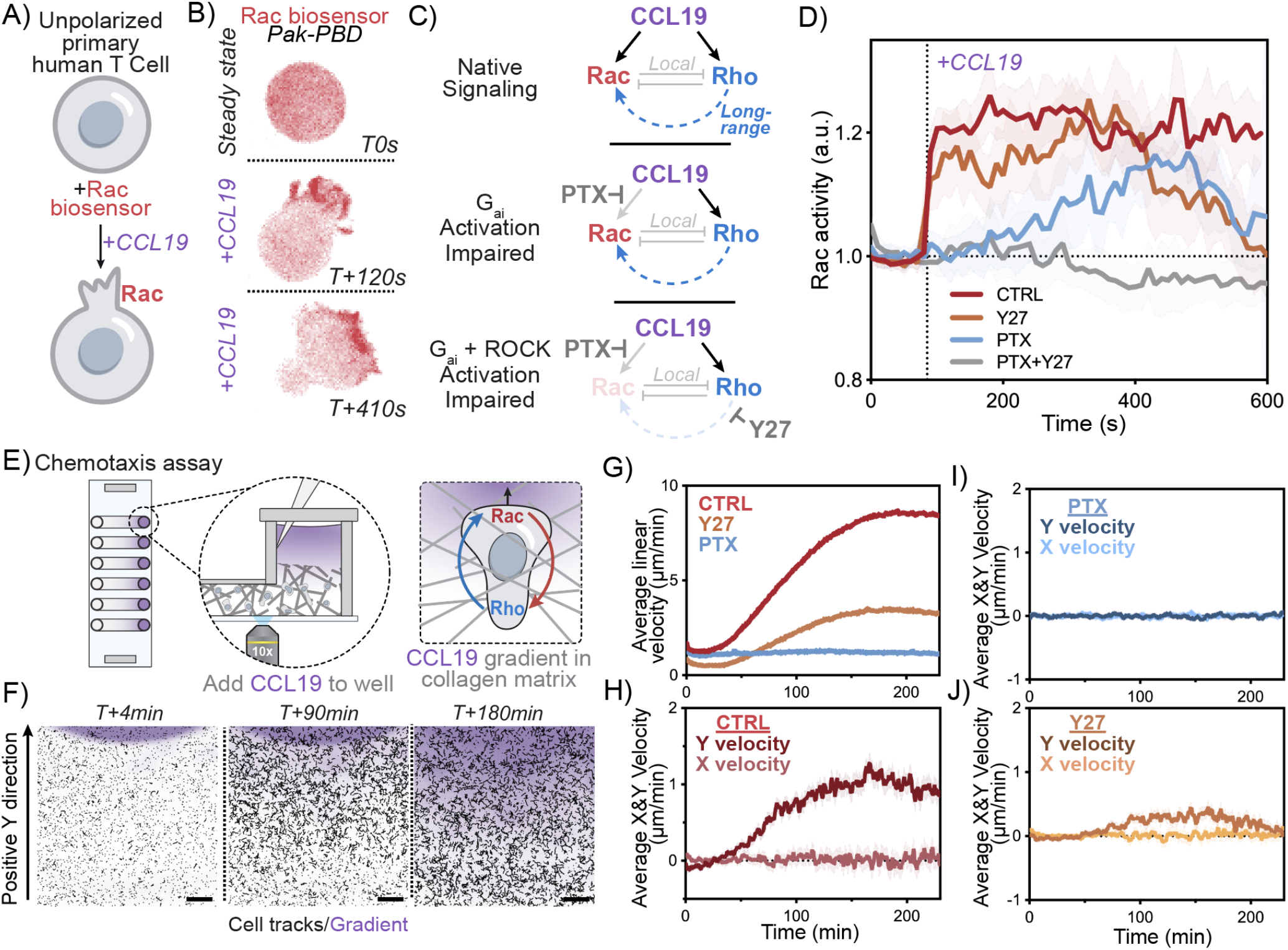
Long range mutual activation establishes robust Rho and Rac polarity during primary T cell migration. (A) Primary human T cells expressing the Rac biosensor (Pak-PBD) to assay cell polarization following acute stimulation with the chemoattractant CCL19. (B) Time-lapse confocal images of an unpolarized human primary T cell before and during CCL19 stimulation. Rac activity was monitored via the Rac biosensor Pak-PBD. (C) Chemoattractants such as CCL19 are known to activate both Rho and Rac. (D) Average time trace of Rac activity at the cell front following addition of 25nM of CCL19 comparing control cells (red), cells treated with 1 *µ*g/ml of G*α*i inhibitor PTX (blue), cells treated with 20*µ*M of Y27 (orange) and a combination of PTX and 20*µ*M Y27 (grey). (mean ± 95%CI; n>20, N = 3). (E) Ex-vivo assay for human primary T cell chemotaxis. Cells are premixed with Bovine Dermal Collagen and placed into linear channels. After collagen sets, media is added to one side of the channel (TCM) and media with human CCL19 (100ng total) and 10ug/mL of Dextran10k-AF647 (Dex647) is added to the other. Imaging takes place right on the edge of the well containing CCL19 so that T cell responses can be recorded as the CCL19 (as read out by Dex647) diffuses into the channel. (F)Representative control experiment is shown at different timepoints with Dex647 fluorescence (top row) and corresponding tracks of T cells (bottom row); tracks display 3 minutes preceding timepoint listed. (G) Average linear velocity of all tracks at each timepoint recorded. Control volunteer N = 3, PTX Volunteer N = 3, Y27 Volunteer N = 2. (H) Average Y only component of velocity of all tracks at each timepoint recorded. Positive is considered towards the well containing CCL19. Control Volunteer N = 3, PTX Volunteer N = 3, Y27 Volunteer N = 2.

### Long range mutual activation establishes robust Rho and Rac polarity during primary T cell migration

Having shown that activating either Rho or Rac leads to a robust establishment of polarity and that Rho and Rac both mutually reinforce each other at a distance, we next investigated the consequences of this long-range cross talk for immune cell polarization and migration. To this end, we used primary human T cells expressing the Rac biosensor (Pak-PBD) to assay cell polarization following acute stimulation with the chemoattractant CCL19 **(Figure 7A)**. We found that CCL19 induces rapid cell polarization with sustained Rac activity at the front **(Figure 7B, S8A-C and Video S9)**.

CCL19 is known to activate both Rac and Rho (50, 51). We first tested if our observed long-lasting Rac partitioning depends solely on the known local activation of Rac via G*α*i signaling or if it also depends on the long-range activation via the Rho/myosin pathway defined in this work **(Figure 7C)**. To focus on the role of Rho/myosin on Rac regulation, we first treated cell with pertussis toxin (PTX) to inactivate G*α*i signaling. These cells maintained their ability to stimulate Rac activation and polarization, albeit with a significant delay and more transiently than control cells **(Figure 7D and S8D-F)**. The ability of PTX treated cells to polarize in response to chemokines has been previously reported but the mechanism by which this is achieved is unknown (6). To test whether this residual Rac regulation occurred through long-range Rho/myosin activation, we treated cells with both PTX and the actomyosin inhibitor Y27632 **(Figure 7D and S8G-I)**. This combination completely abolished even transient Rac polarization, consistent with a long-range back-to-front signaling link that acts in conjunction with G*α*i-based Rac activation.

We previously showed that long-range Rac activation by Rho is dependent on PI3K which locally converts PIP2 into the Rac activator PIP3 at the cell front. We further investigated whether chemoattractant induced back-to-front signaling is also dependent on PI3K activation and found Rac polarization to be blocked when using both PTX (which blocks the Gai-based Rac activation) and the PI3K inhibitor Duvelisib (which blocks the long-range Rho-based Rac activation, **Figure S8J-L**). Finally, we show that preventing the long-range Rho to Rac communication using Y27632 alone elicits a more transient polarization than is seen for control cells **(Figure S8M-O)**. These results demonstrate that chemoattractant signaling communicates with the front program in two different ways, one direct route via G*α*i and one indirect route via the back-mediated mechanism we define in this work. Cells deficient in either the direct or indirect(/long-range) route can only polarize transiently, while WT primary cells combine both direct and indirect routes of Rac activation for robust and sustained polarization in response to chemoattractant.

Finally, we sought to test the role of this robust polarization established by the long-range Rac-Rho communication during chemoattractant induced migration. To stimulate chemotaxis, we embedded T cells in a 3D collagen matrix in a thin channel with CCL19 placed on one side **(Figure 7E)**. As CCL19 freely diffuses into the channel, a chemotactic gradient is established, and control cells rapidly polarize and chemotax up the CCL19 gradient **(Figure 7F-H and Video S10)**. In contrast, cells treated with PTX or Y27 only briefly polarized and failed to robustly migrate toward the CCL19 gradient **(Figure 7G, I-J and S8Q)**.

Our results show that long-range mutual activation of front and back is an essential feature of human lymphocyte polarization and motility. Furthermore, we show that activating either the front or back polarity programs without the other only leads to transient polarization. Dual activation is required to support sustained polarization and efficient chemotactic migration.

## Discussion

Our work demonstrates that the well-appreciated mutual short-range inhibition between Rac and Rho is accompanied by an additional mutual long-range facilitation of the front and back polarity programs in migrating cells **(Figure 8)**. The front reinforces the back via membrane tension mediated mTORC2 activation **(Figure 1, 2 S1-4)** The back reinforces the front via cortical remodeling and localized PIP2 release **(Figure 3, 4 and S5, S6)**. Both modes of long-range facilitation are required for sustained and efficient front organization and chemotaxis in primary human lymphocytes **(Figure 7)**. Understanding the partitioning of Rho and Rac in migrating cells has long been a challenge due to the difficulty in establishing the range and directionality of interactions of Rac and Rho. Our use of optogenetics in unpolarized cells enables us to dissect the long-range positive interactions between the front and back programs, and our combined mechanical and biochemical models complement our experimental work by revealing the importance of these links to cell polarization. Our combined multidisciplinary approach is likely to be similarly powerful for other complex interconnected spatiotemporal signaling networks such as axon specification (52), asymmetric cell division (53), epithelial polarity (54), planar cell polarity (55), branching morphogenesis (56), lumen formation (57), and collective cell migration (58).

**Fig. 8.**
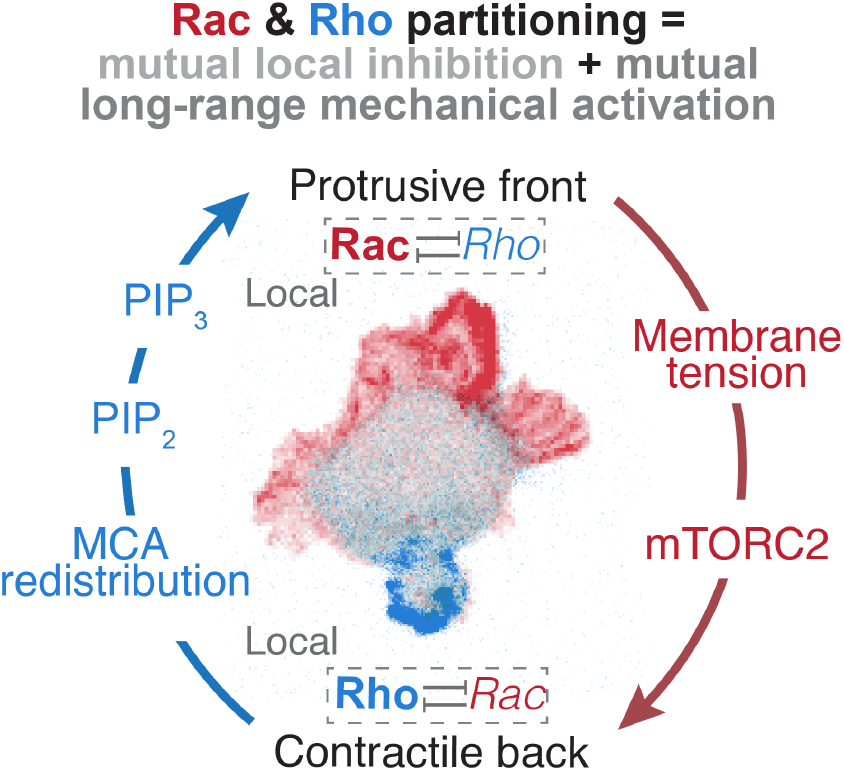
Summary figure.

Robust cell polarization necessitates long-range integration of signals across the cell. A number of currencies have been proposed for this long-range signal relay: diffusion of signals across the cell (14, 17, 47, 59, 60) or force-mediated communication via propagation of forces in the membrane or cortex (26–28, 30, 61) or advection of cellular components via force asymmetries across the cell (62–64). Membrane tension rapidly propagates across cells and is used for long-range integration of cellular processes from cell spreading (65, 66) to phagocytosis (67) to the competition of fronts that enables a dominant protrusion in migrating cells (27, 28). Tension-based communication is particularly powerful for global signal transmission across the cell (37). Our work shows that membrane tension also enables the front to facilitate contraction at the cell rear through mTORC2 activation. Previous studies (34, 68–73), including our own (28, 29), have broadly linked TORC2 and mTORC2 to cell polarity, but how mTORC2 does so is not well-understood. Even its links to myosin regulation have been a point of contention, with some studies showing a positive role (29) and others proposing a negative role (34). Our work provides direct evidence for and a molecular mechanism for this link **(Figure 2)**. The evolutionarily-conserved mTORC2 is also the mechanosensitive pathway that enables competition between fronts (28, 29), regulation of plasma membrane homeostasis in budding yeast (74, 75), and chemotaxis in cells from neutrophils (34) to Dictyostelium (70). Given the evolutionarily conserved role of mTORC2, it would be interesting to probe in future work whether the mTORC2 mediated activation of Rho in response to elevated membrane tension is important in other contexts such as cell division.

Force-mediated flow of components is another powerful mechanism of long-range cell communication (62, 76, 77). Our work shows that flow of MCA components secondary to Rho-mediated actin-based contractions enables the cell rear to establish a Rac-based protrusive front at the opposite end of the cell. These flow-mediated modes of communication are an efficient means for the polarized distribution of signals—in this case the depletion of MCA sites and the facilitation of Rac activation at the opposite end of the cell from Rho-mediated contractions. Flow of cellular signals from the cell front to the cell back along retrograde flows or cortical contractions have been proposed to enable robust polarization (62, 76, 77). Our work provides a molecular mechanism for this long-range integration. It is interesting that the long-range back-to-front and front-to-back communication involve different currencies of membrane tension and cortical flow. These different currencies have different spatial patterns of propagation (relatively global for membrane tension and polarized for cortical flows) and also differ in their molecular requirements, potentially enabling these modes of communication to be independently modulated without interfering with one another.

Rho GTPases are central regulators of the cytoskeleton, and their partitioning underlie many fundamental cellular functions such as cell shape, adhesions and division1. In the context of cell locomotion, we find that long-range activation is a key complement to local inhibition to ensure the proper partitioning of Rac and Rho at opposite poles of the cell. Our findings and model highlight the limitations of a system that relies only on local interactions for spatial segregation. We speculate that the coupling between local biochemical interactions and long-range mechanical facilitation is a conserved feature of Rho GTPases partitioning. This is in good agreement with other systems in which Rho GTPase partitioning relies on long-range communication via the actin and the membrane, from polarization in early C. elegans embryos by cortical flow (78), to axon branching in neuron via long-range membrane tension propagation (52), to the winner-take-all competition enabling a single front during migration (27, 28). Long-range mechanochemical interactions represent a conserved mechanism to complement local biochemical interactions for the partitioning of Rho GTPases in cell migration and likely other processes.

## ACKNOWLEDGEMENTS

We thank the Weiner and Turlier lab for their critical feedback. We especially want to thank Kate Cavanaugh for kindly sharing the PIP2 biosensor construct and Juan Manuel García-Arcos for his helpful insights. We also thank the blood donors that made the primary human T cell experiment possible. This work was supported by R35GM118167 (ODW, ES), National Science Foundation/Biotechnology and Biological Sciences Research Council grant 2019598 (ODW), K99GM154115 (HDB), the European Research Council (ERC) under the European Union’s Horizon 2020 research and innovation program (Grant agreement No. 949267) (HT), the European Union’s Horizon 2020 research and innovation program under the Marie Sklodowska-Curie grant agreement No. 101150259 (AFG), the American Heart Association predoctoral fellowship 24PRE1198865 (PJZ) and R21AI1739938 (JKB). HT and AFG are supported by the CNRS and Collège de France.

