## Supplementary Materials for "Long range mutual activation establishes Rho and Rac polarity during cell migration"

**Supplementary figure and videos legends:**

**Video S1**

Time-lapse confocal images of HL-60 cells expressing opto-PI3k (Opto-Rac) and Rac biosensor (Pak-PBD) showing localized Rac activation upon light activation. Related to **Figure 1C, D**. Scale bar: 10um.

**Video S2**

Time-lapse confocal images of HL-60 cells expressing opto-PI3k (Opto-Rac) and Rho biosensor (AnillinRBD) showing that light-induced Rac activation elicits a rapid long-range increase in Rho activity at the opposite side of the cell. Related to **Figure 1F, G**. Scale bar: 10um.

**Video S3**

Time-lapse confocal images of HL-60 cells expressing Rho biosensor (AnillinRBD) before and during osmotic shock performed by directly adding water to the imaging media (70% final). This shows that Rho activity increase in response to increasing membrane tension by osmotic shock. Related to **Figure 2G, H**. Scale bar: 10um.

**Video S4**

Time-lapse confocal images of an HL-60 cell expressing Rho biosensor (AnillinRBD) and stained with membrane marker CellMask before and during micropipette aspiration assay, showing that Rho activity increase in response to mechanically increasing membrane tension. Related to **Figure 2K, L**. Scale bar: 10um.

**Video S5**

First video: Time-lapse confocal images of a Rictor KO HL-60 cell expressing opto-PI3k (Opto-Rac) and Rho biosensor (AnillinRBD) showing that light-induced Rac activation elicits no significant increase in Rho activity at the opposite side of the cell. Related to **Figure 3B, C**. Scale bar: 10um.

Second video: Time-lapse confocal images of a Rictor KO HL-60 cell expressing Rho biosensor (AnillinRBD) before and during osmotic shock performed by directly adding water to the imaging media (70% final) showing no significant change in Rho activity in response to osmotic shock. Related to **Figure 3E, F**. Scale bar: 10um.

Third video: Time-lapse confocal images of a Rictor KO HL-60 cells expressing Rho biosensor (AnillinRBD) and stained with membrane marker CellMask before and during micropipette aspiration assay, showing no significant Rho activity increase in response to mechanically increasing membrane tension. Related to **Figure 3H, I**. Scale bar: 10um.

**Video S6**

Time-lapse confocal images of an HL-60 cell expressing opto-LARG (Opto-Rho) and the Rac biosensor (Pak-PBD) showing that Rho activation leads to long-range Rac activation at the opposite end of the cell, coinciding with another morphological change— blebbing. Related to **Figure 4C, D**. Scale bar: 10um.

**Video S7**
Time-lapse confocal images of an HL-60 cell expressing opto-LARG (Opto-Rho) and Ezrin-mCherry showing that upon local Rho activation, ezrin accumulated at the site of contraction and was depleted at the opposite side of the cell Related to **Figure 5B, C**. Scale bar: 10um.

**Video S8**
Time-lapse confocal images of an HL-60 cell expressing opto-LARG (Opto-Rho) and PIP2 biosensor (Tubby-HaloTag) showing that local Rho activation leads to a marked increase in $PIP_2$ level in blebs. Related to **Figure 5F, G**. Scale bar: 10um.

**Video S9**
Time-lapse confocal images of a primary T Cell expressing the Rac biosensor (Pak-PBD) before and after adding 25nM of CCL19. Related to **Figure 7C, D**.

**Video S10**
Ex-vivo assay for human primary T cell chemotaxis. Cells are premixed with Bovine Dermal Collagen and placed into linear channels. After collagen sets, media is added to one side of the channel (TCM) and media with human CCL19 (100ng total) and 10ug/mL of Dextran10k-AF647 (Dex647) is added to the other. Average linear velocity of all tracks at each timepoint is recorded. In order showing control cells, cells treated with Y27 and cells treated with PTX. Related to **Figure 7E-J**.

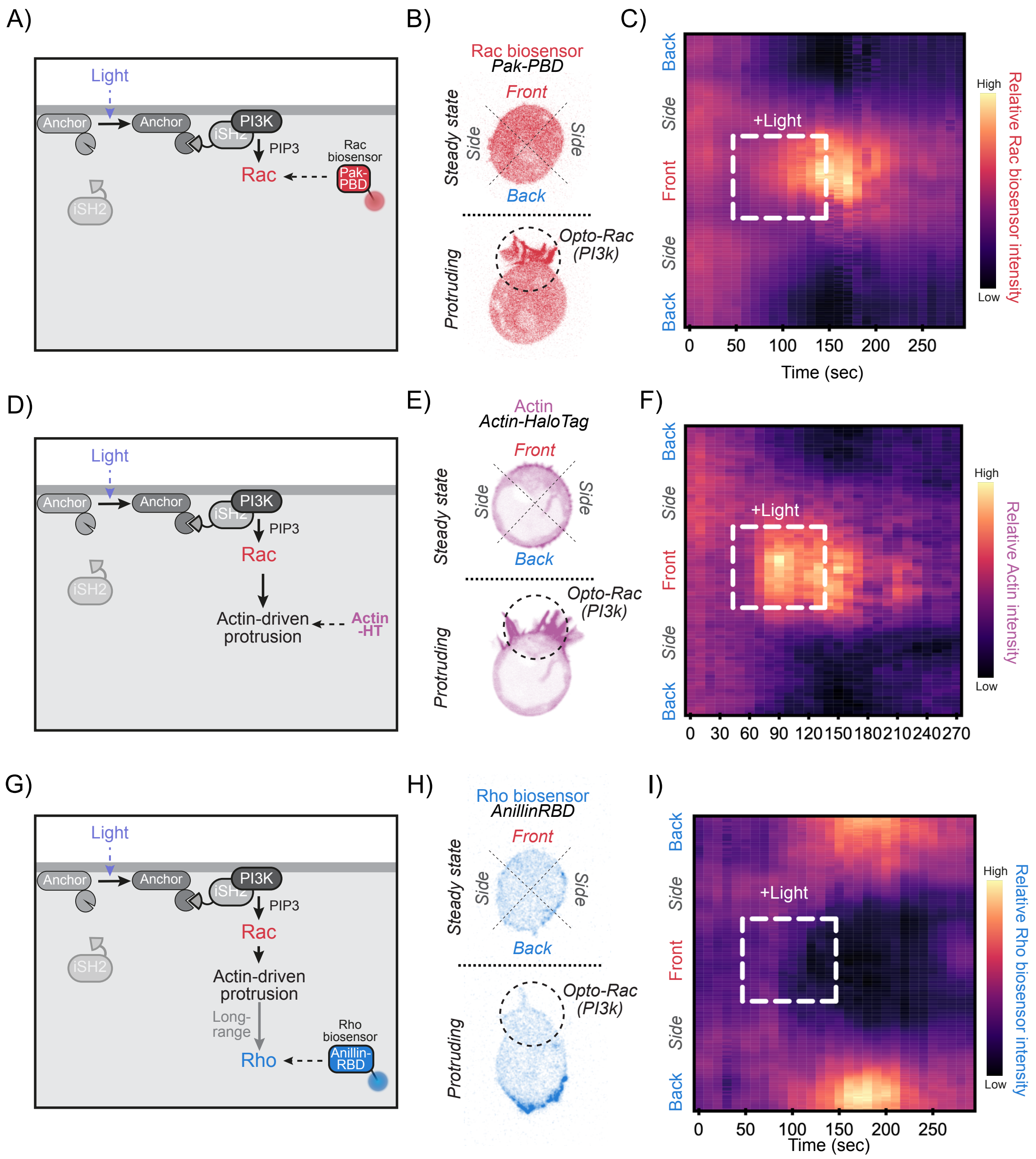
A)
Light
Anchor
Anchor
PI3K
iSH2
PIP3
Rac
Rac biosensor
Pak-PBD
iSH2
B)
Rac biosensor
Pak-PBD
Front
Side
Side
Back
Steady state
Opto-Rac (PI3k)
Protruding
C)
Back
Side
Front
Side
Back
+Light
High
Low
Relative Rac biosensor intensity
0 50 100 150 200 250
Time (sec)
D)
Light
Anchor
Anchor
PI3K
iSH2
PIP3
Rac
Actin-driven protrusion
Actin -HT
iSH2
E)
Actin
Actin-HaloTag
Front
Side
Side
Back
Steady state
Opto-Rac (PI3k)
Protruding
F)
Back
Side
Front
Side
Back
+Light
High
Low
Relative Actin intensity
0 30 60 90 120 150 180 210 240 270
G)
Light
Anchor
Anchor
PI3K
iSH2
PIP3
Rac
Actin-driven protrusion
Long-range
Rho
Rho biosensor
Anillin-RBD
iSH2
H)
Rho biosensor
AnillinRBD
Front
Side
Side
Back
Steady state
Opto-Rac (PI3k)
Protruding
I)
Back
Side
Front
Side
Back
+Light
High
Low
Relative Rho biosensor intensity
0 50 100 150 200 250
Time (sec)

**Figure S1: Optogenetic control of PI3K leads to local Rac activation, which triggers localized actin-driven cell protrusion and long-range Rho activation.**

(A) Schematic of local Rac activation via opto-PI3K. When localized 488-nm light is applied, the membrane anchor protein (iLid-BFP-CAAX) changes conformation, which leads to the binding of inter SH2 domain (iSH2) to the illuminated region. iSH2 then recruits PI3K, whose lipid product ($PIP_3$) induces the activation of Rac GTPase (Rac). We monitor Rac activity using the Rac biosensor Pak-PBD-mCherry, which recognizes and binds the active GTP-bound Rac. (B) Time-lapse confocal images of an unpolarized cell before and during opto-Rac stimulation. Rac activity was monitored via the Rac biosensor Pak-PBD. (C) Kymograph of Rac activity (Pak-PBD intensity) along the normalized cell circumference (y axis) showing that over time (x axis), Rac activity increases at the front in response to opto-PI3K. (N=3, n>40). (D) Rac activation via opto-PI3K leads to localized actin-mediated protrusion, which we monitor by over-expression of Actin-HaloTag. (E) Time-lapse confocal images of an unpolarized cell expressing actin-HaloTag before and during opto-Rac stimulation. (F) Kymograph of actin intensity during opto-PI3k activation (N = 3, n = 25). (G) Long-range Rho activation by local Rac activation is monitored using the Rho Biosensor AnillinRBD, which recognizes the active GTP-bound Rho. (H) Time-lapse confocal images of an unpolarized cell expressing AnillinRBD-mCherry before and during opto-Rac stimulation. (I) Kymograph of Rho activity in response to local Rac activation (opto-PI3k) (N = 8, n > 40).

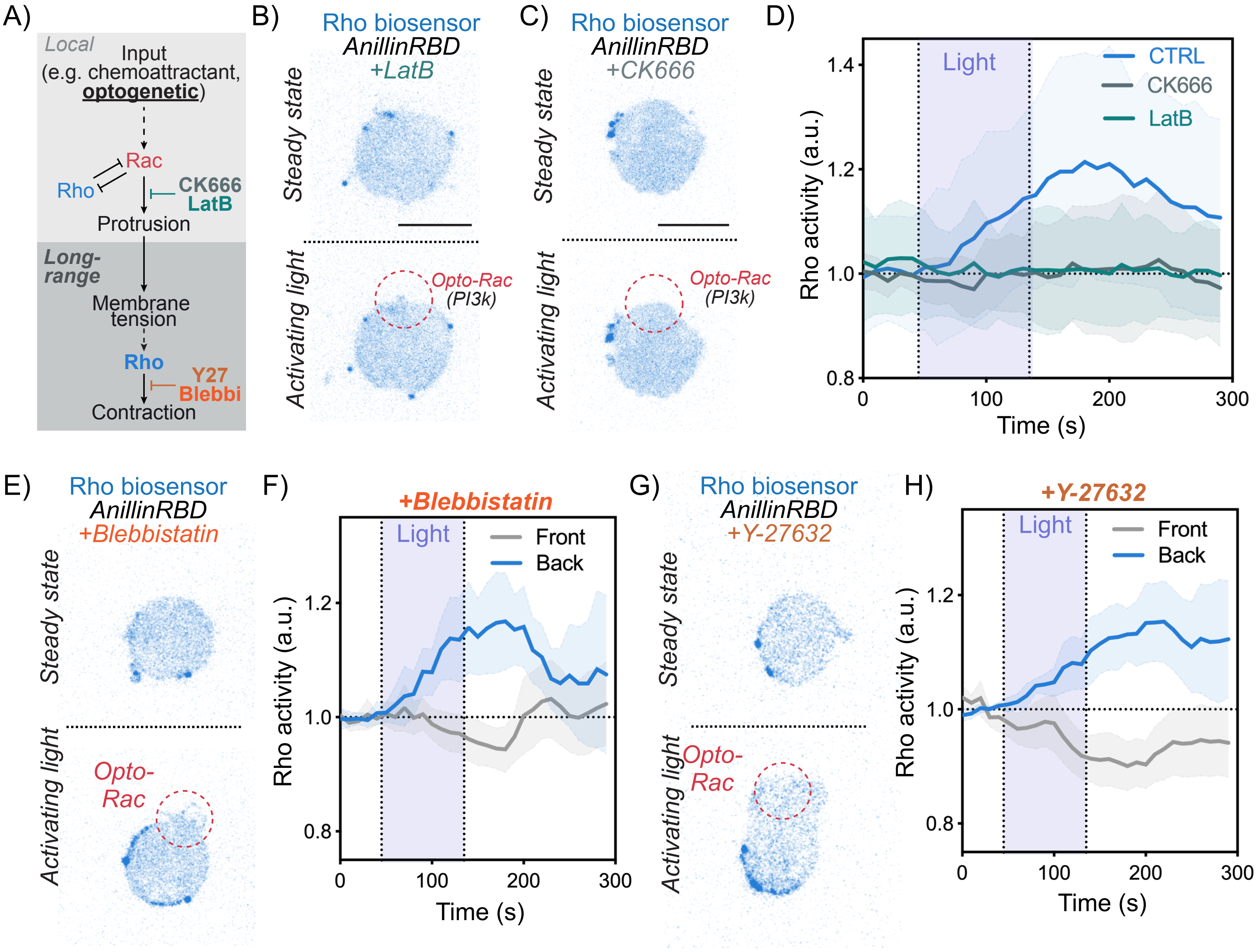
A)
Local
Input
(e.g. chemoattractant, optogenetic)
Rac
Rho
CK666
LatB
Protrusion
Long-range
Membrane tension
Rho
Y27
Blebbi
Contraction
B)
Rho biosensor
AnillinRBD
+LatB
Steady state
Activating light
Opto-Rac
(PI3k)
C)
Rho biosensor
AnillinRBD
+CK666
Steady state
Activating light
Opto-Rac
(PI3k)
D)
Light
CTRL
CK666
LatB
Rho activity (a.u.)
Time (s)
E)
Rho biosensor
AnillinRBD
+Blebbistatin
Steady state
Activating light
Opto-Rac
F)
+Blebbistatin
Light
Front
Back
Rho activity (a.u.)
Time (s)
G)
Rho biosensor
AnillinRBD
+Y-27632
Steady state
Activating light
Opto-Rac
H)
+Y-27632
Light
Front
Back
Rho activity (a.u.)
Time (s)

**Figure S2: Long-range Rho activation requires cellular protrusion but not contraction**

(A) To test the role of protrusions in Rac/Rho crosstalk, we locally activated Rac via optogenetics in the presence of inhibitors of actin assembly (Latrunculin B) or the Arp2/3 complex activation (CK666). (B)(C) Time-lapse confocal images of unpolarized cells expressing AnillinRBD-mCherry before and during opto-Rac stimulation treated with either 10 $\mu$M of Latrunculin B (B) or 100 $\mu$M of CK666 (C).(D) Average time trace of Rho activity at the plasma membrane at the back of cells treated with either 10 $\mu$M of Latrunculin B (teal curve) or 100 $\mu$M of CK666 (green curve) demonstrate minimal change in Rho activity following opto-Rac activation (mean $\pm$ 95%CI; n>30, N= 3) in contrast to control cells in which the cytoskeleton was not inhibited (same curve as in **Fig. 1D**). (E) Confocal images of opto-PI3K cells expressing the Rho biosensor AnillinRBD treated with 10$\mu$M of para-amino-blebbistatin (Myosin inhibitor) before and during light-induced protrusions. (F) Average time trace of Rho activity at the plasma membrane at the back of cells treated with 10 $\mu$M of para-amino-blebbistatin demonstrate that myosin contractility is not required for the protrusion-mediated long-range Rho activation (mean $\pm$ 95%CI; n = 20, N= 2) (G) (H) Same as (E)(F) but using cells treated with 20$\mu$M of the ROCK inhibitor Y-27632, showing that contractility is not required for the protrusion-mediated long-range Rho activation (mean $\pm$ 95%CI; n = 20, N= 2).

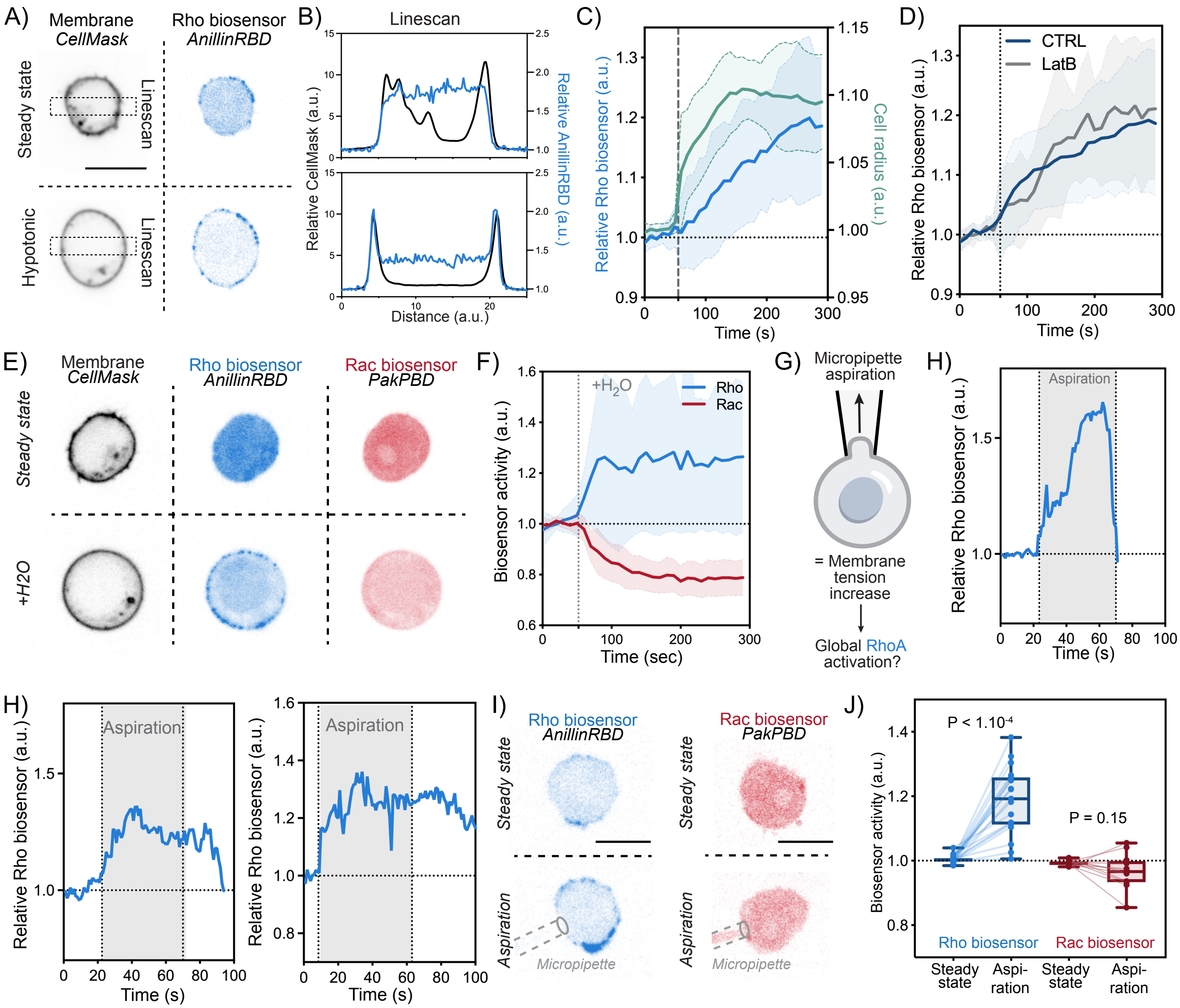

A)
Membrane
CellMask
Rho biosensor
AnillinRBD
Steady state
Linescan
Hypotonic
Linescan
B)
Linescan
Relative CellMask (a.u.)
Relative AnillinRBD (a.u.)
Distance (a.u.)
C)
Relative Rho biosensor (a.u.)
Cell radius (a.u.)
Time (s)
D)
CTRL
LatB
Relative Rho biosensor (a.u.)
Time (s)
E)
Membrane
CellMask
Rho biosensor
AnillinRBD
Rac biosensor
PakPBD
Steady state
+H2O
F)
+H2O
Rho
Rac
Biosensor activity (a.u.)
Time (sec)
G)
Micropipette aspiration
= Membrane tension increase
Global RhoA activation?
H)
Aspiration
Relative Rho biosensor (a.u.)
Time (s)
H)
Aspiration
Relative Rho biosensor (a.u.)
Time (s)
Aspiration
Relative Rho biosensor (a.u.)
Time (s)
I)
Rho biosensor
AnillinRBD
Rac biosensor
PakPBD
Steady state
Aspiration
Micropipette
J)
P < 1.10-4
P = 0.15
Biosensor activity (a.u.)
Rho biosensor
Rac biosensor
Steady state
Aspi-ration
Steady state
Aspi-ration

**Figure S3: Rho, unlike Rac, is activated by increased membrane tension**

(A) Confocal images of a cell before and during osmotic shock (+70% water) expressing the Rho biosensor AnillinRBD and stained with the plasma membrane marker CellMask. (B) linescan of the Rho and membrane intensity before and during osmotic shock, showing a decrease in background biosensor in the cytoplasm (cytoplasmic dilution) and a recruitment of the biosensor at the plasma membrane. (C) Left y axis, average time trace of Rho activity at the membrane (in blue) in response to hypotonic shock (70% water). Right y axis, cell radius over time (green). (N = 2, n >15, means ± 95%CI) (D) Average time trace of Rho activity at the membrane in response to hypotonic shock (80% water) for control cells (blue) or cells treated with 10 uM of Latrunculin B (grey), showing that actin inhibition doesn't impair Rho recruitment in response to elevated membrane tension. (N = 3, n >30, means ± 95%CI). (E) Confocal images of a cell dyed with the membrane marker CellMask and expressing both Rho (Anillin-RBD) and Rac (Pak-BPD) biosensors before and during osmotic shock (+70% water) (F) Average time trace of Rho and Rac activity at the plasma before and after hypo-osmotic shock (60 mOsm or 70% H2O). Hypotonic-shock-based elevation of membrane tension suffices to globally increase RhoA activity and leads to a global decrease in Rac activity as was previously reported[28] (mean ± 95%CI; n>25, N= 3). (G) As an alternate approach to increase membrane tension, we leveraged micropipette aspiration[32] (H) Three example time trace of Rho activity at the plasma membrane in response to micropipette aspiration. (I) Time-lapse confocal images of an unpolarized opto-Rac HL-60 cell expressing either a Rho biosensor (AnillinRBD) or a Rac biosensor (PakPBD) before and after micropipette aspiration. (J) Average Rho and Rac activity before (steady state) and during aspiration. Micropipette-based elevation of membrane tension significantly stimulates Rho activation while leading to a non-significant change in Rac activity. Box and whiskers: median and min to max; p values from Wilcoxon paired Student's t test. Scale bars: 10μm.

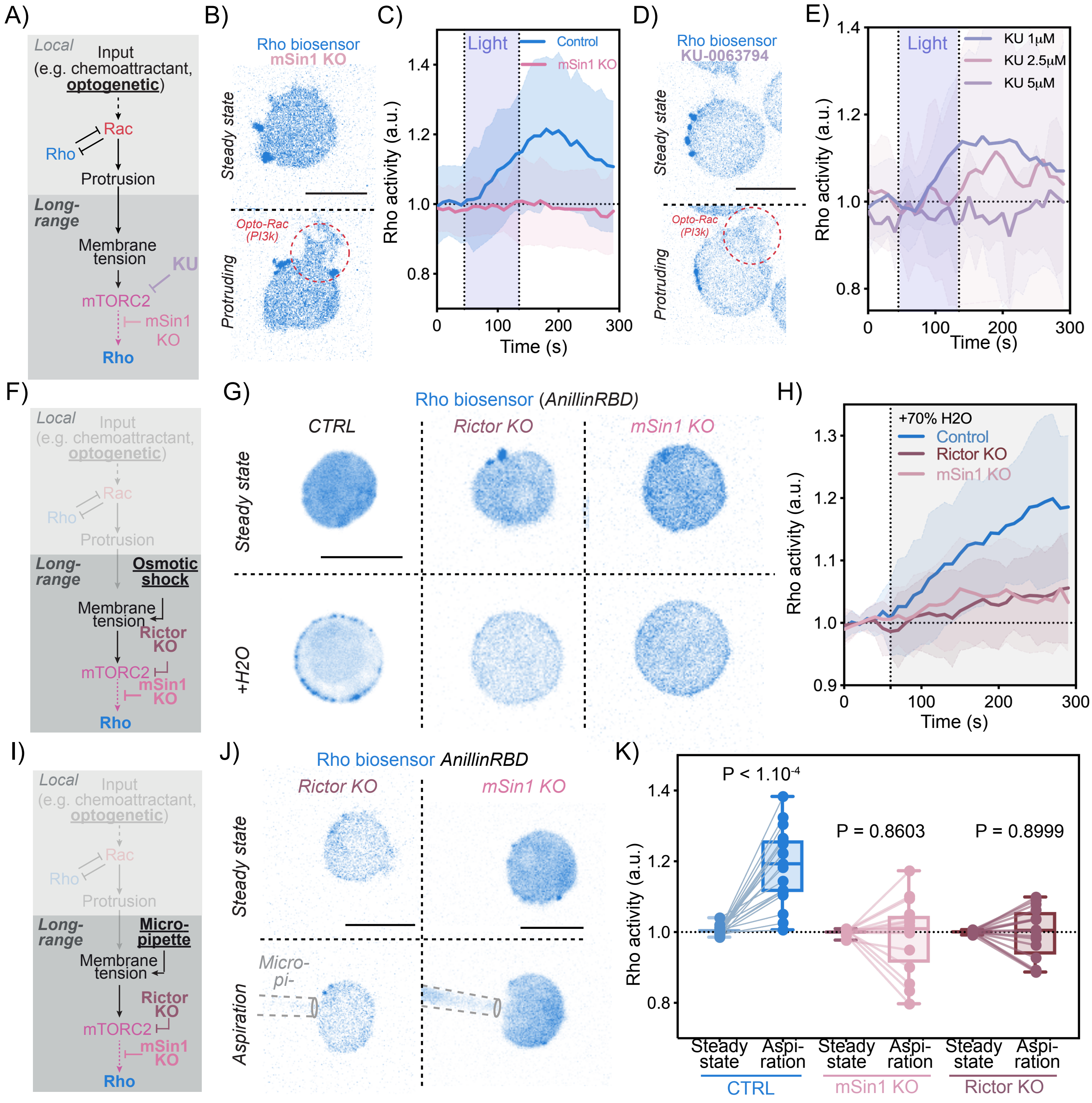

A)
Local
Input
(e.g. chemoattractant, optogenetic)
Rac
Rho
Protrusion
Long-range
Membrane tension
KU
mTORC2
mSin1 KO
Rho
B)
Rho biosensor
mSin1 KO
Steady state
Opto-Rac (PI3k)
Protruding
C)
Light
Control
mSin1 KO
Rho activity (a.u.)
Time (s)
D)
Rho biosensor
KU-0063794
Steady state
Opto-Rac (PI3k)
Protruding
E)
Light
KU 1μM
KU 2.5μM
KU 5μM
Rho activity (a.u.)
Time (s)
F)
Local
Input
(e.g. chemoattractant, optogenetic)
Rac
Rho
Protrusion
Long-range
Osmotic shock
Membrane tension
Rictor KO
mTORC2
mSin1 KO
Rho
G)
Rho biosensor (AnillinRBD)
CTRL
Rictor KO
mSin1 KO
Steady state
+H2O
H)
+70% H2O
Control
Rictor KO
mSin1 KO
Rho activity (a.u.)
Time (s)
I)
Local
Input
(e.g. chemoattractant, optogenetic)
Rac
Rho
Protrusion
Long-range
Micro-pipette
Membrane tension
Rictor KO
mTORC2
mSin1 KO
Rho
J)
Rho biosensor AnillinRBD
Rictor KO
mSin1 KO
Steady state
Micro-pi-
Aspiration
K)
P < 1.10-4
P = 0.8603
P = 0.8999
Rho activity (a.u.)
Steady state
Aspi-ration
CTRL
mSin1 KO
Rictor KO

**Figure S4: mTORC2 links membrane tension increases to Rho activation**

We investigate whether mTORC2 is part of the mechanosensory pathway that links increased in membrane tension to the activation of Rho. (A) We used optogenetics to locally activate Rac in control cells versus cells with impaired mTORC2 activity (either by using cells lacking mSIN1, a core component of the mTORC2 complex or cells treated with the mTOR inhibitor KU-0063794. (B) Time-lapse confocal images of an unpolarized control or mSIN1 KO cell before and during opto-Rac stimulation. Rac activity was monitored via the Rac biosensor Pak-PBD. (C) Local Rac activation potently stimulates long-range activation of Rho in control cells (blue) but not cells deficient in mTORC2 activity (mSIN1 KO, pink) (mean ± 95%CI; n>20, N= 3). Control curve is same as in **Fig. 1D**. (D) Time-lapse confocal images of unpolarized cells expressing AnillinRBD-mCherry treated with 5uM of KU-0063794 before and during opto-Rac stimulation. (E) Average time trace of Rho activity in opto-Rac cells treated with either 1uM (blue), 2.5uM (pink) or 5uM (purple) of KU-0063794. (N = 2, n > 8). (F) Osmotic shock to globally increase membrane tension in cells lacking core components of the mTORC2 complex (either Rictor or mSIN1 knockouts). (G) Time-lapse confocal images of unpolarized control, Rictor KO or mSIN1 KO cells expressing AnillinRBD-mCherry before and during hypotonic shock (70% Water). (H) Average time trace of Rho activity at the plasma membrane before and after hypo-osmotic shock in control, Rictor KO and mSIN1 KO cells demonstrate a requirement for mTORC2 in membrane-tension mediated activation of RhoA (mean ± 95%CI; n>35, N= 3). (I) Same as (F) but using micropipette aspiration to mechanically stimulate an increase in membrane tension. (J) Time-lapse confocal images of unpolarized Rictor KO or mSIN1 KO HL60s cells expressing the Rho biosensor (AnillinRBD) before and after micropipette aspiration. (K) Average Rho activity before (steady state) and during aspiration of control, Rictor KO and mSIN1 KO cells. Values for control are the same as **Fig. 2G**. Box and whiskers: median and min to max; p values from Wilcoxon paired Student's t test. Scale bars: 10μm.

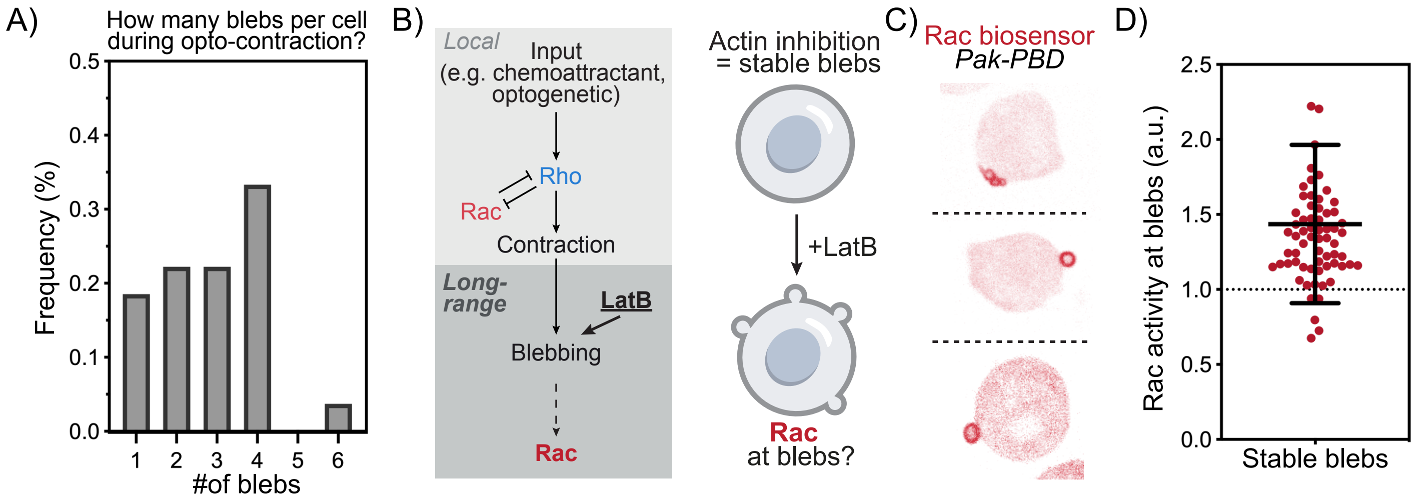
A)
How many blebs per cell during opto-contraction?
Frequency (%)
0.5
0.4
0.3
0.2
0.1
0.0
1 2 3 4 5 6
#of blebs
B)
Local
Input
(e.g. chemoattractant, optogenetic)
Rho
Rac
Contraction
Long-range
LatB
Blebbing
Rac
Actin inhibition = stable blebs
+LatB
Rac at blebs?
C)
Rac biosensor
Pak-PBD
D)
Rac activity at blebs (a.u.)
2.5
2.0
1.5
1.0
0.5
0.0
Stable blebs

**Figure S5: Blebs are sufficient for Rac activation**

(A) Histogram of the number of blebs per cell in response to opto-Rho stimulation (N = 3, n>55). (B) To determine whether blebs suffice for Rac activation in a context independent of Rho-mediated stimulation of actomyosin contractility, we used the actin inhibitor Latrunculin B to generate stable blebs. (C) Three representative confocal images of cells expressing the Rac biosensor Pak-PBD treated with 10 μM of Latrunculin B demonstrate Rac enrichment in stable blebs. (D) Quantitation of Rac activity in LatrunculinB-induced stable blebs (see Methods) (mean ± 95%CI; n>70, N= 3).

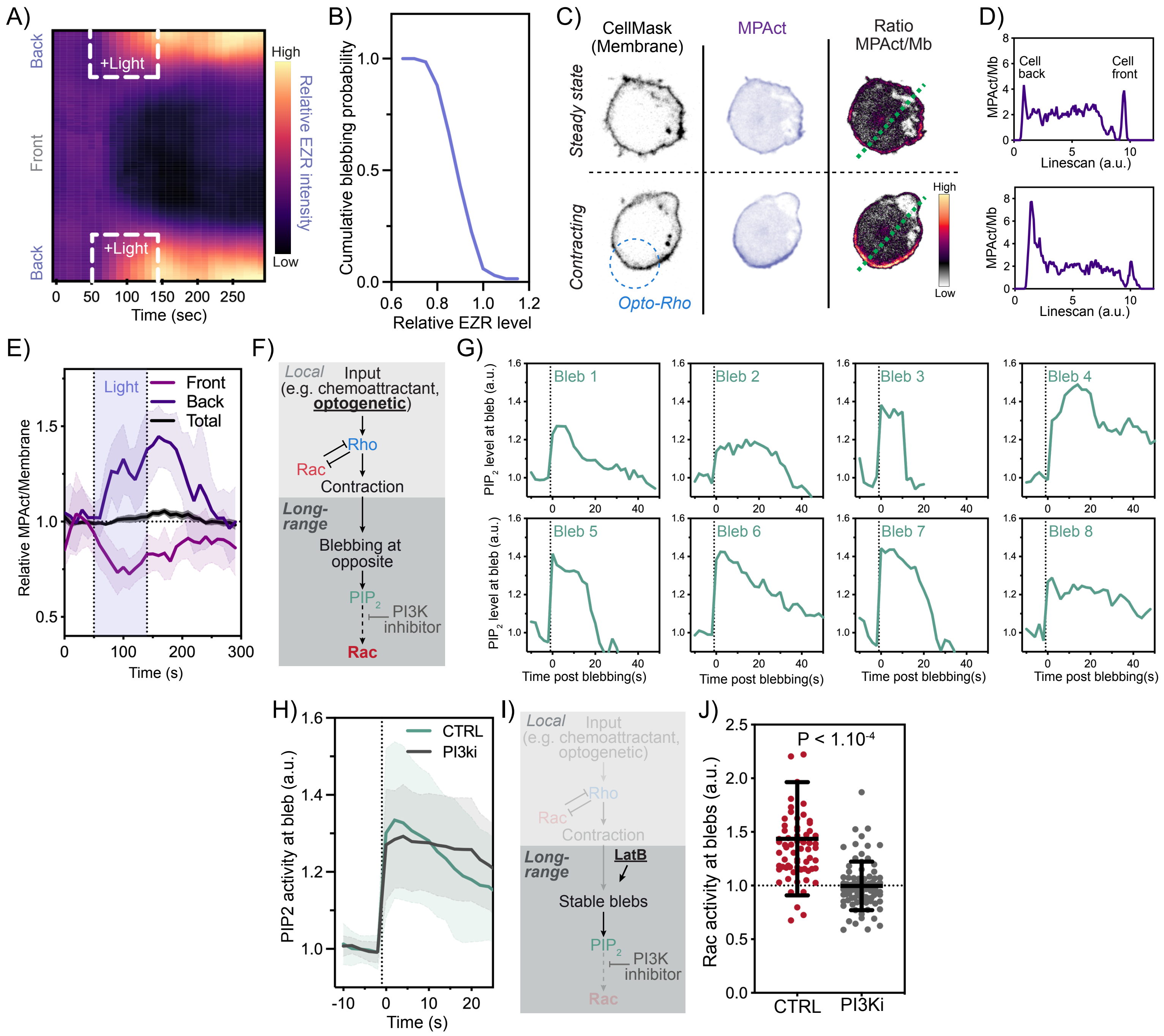

A)
Back
Front
Back
+Light
+Light
High
Low
Relative EZR intensity
Time (sec)
B)
Cumulative blebbing probability
Relative EZR level
C)
CellMask (Membrane)
MPAct
Ratio MPAct/Mb
Steady state
Contracting
Opto-Rho
High
Low
D)
Cell back
Cell front
MPAct/Mb
Linescan (a.u.)
E)
Light
Front
Back
Total
Relative MPAct/Membrane
Time (s)
F)
Local
Input (e.g. chemoattractant, optogenetic)
Rho
Rac
Contraction
Long-range
Blebbing at opposite
PIP2
PI3K inhibitor
Rac
G)
Bleb 1
Bleb 2
Bleb 3
Bleb 4
Bleb 5
Bleb 6
Bleb 7
Bleb 8
PIP2 level at bleb (a.u.)
Time post blebbing(s)
H)
CTRL
PI3ki
PIP2 activity at bleb (a.u.)
Time (s)
I)
Local
Input (e.g. chemoattractant, optogenetic)
Rho
Rac
Contraction
Long-range
LatB
Stable blebs
PIP2
PI3K inhibitor
Rac
J)
P < 1.10-4
Rac activity at blebs (a.u.)
CTRL
PI3Ki

**Figure S6: Contraction-induced membrane-to-cortex attachment asymmetry leads to $PIP_2$ release and PI3K-dependent Rac activation at the opposite side of the cell**

(A) In unpolarized cells at steady state, actin and MCA proteins are uniformly distributed. Upon local contraction (for example induced by opto-Rho), Actin flows toward the site of contraction. We ask whether MCA flows with the cytoskeleton toward the site of contraction to generate an MCA asymmetry across the cell (B) Kymograph of Ezrin intensity along the normalized cell circumference (y axis) showing that over time (x axis), Ezrin is enriched at the back in response to opto-Rho activation. (C) Cumulative blebbing probability in function of relative Ezrin level. (N = 3, n >40). (D) Time-lapse confocal images of unpolarized cells expressing the MPact, a membrane-actin proximal sensor[43] which we use as a proxy for MCA level, and stained with the membrane marker CellMask before and during opto-Rho stimulation. The ratio between MPact and membrane channel is used to measure the approximative MCA level intensity. (E) Line-scan of MPAct/Membrane ratio of cell displayed in (D), see green dotted line. (F) Average time trace of relative MPAct/Membrane ratio (normalized to before light) at the front and back of the cell. (N = 2, n >10, means ± 95%CI). (G) Local Rho-induced contractions lead to MCA flow toward the site of contraction and depletion of MCA at the opposite end of the cell, resulting in blebbing at the cell pole that is opposed to opto-Rho. We hypothesized that MCA detachment from plasma membrane during blebbing leads to $PIP_2$ release. (H) Average time trace of $PIP_2$ activity at the bleb in control cells and cells treated with 1 uM PI3Kδ/γ inhibitor. (N = 3, n >30, means ± 95%CI) (I) To determine whether blebs suffice for $PIP_2$-mediated Rac activation in a context independent of Rho-mediated stimulation of actomyosin contractility, we used the actin inhibitor Latrunculin B to generate stable blebs. (J) Rac activity in stable blebs in control cells and cells treated with 1 uM PI3Kδ/γ inhibitor show that PI3K inhibition prevents Rac activation in stable blebs. (N = 3, n >70, p values from Mann Whitney test).

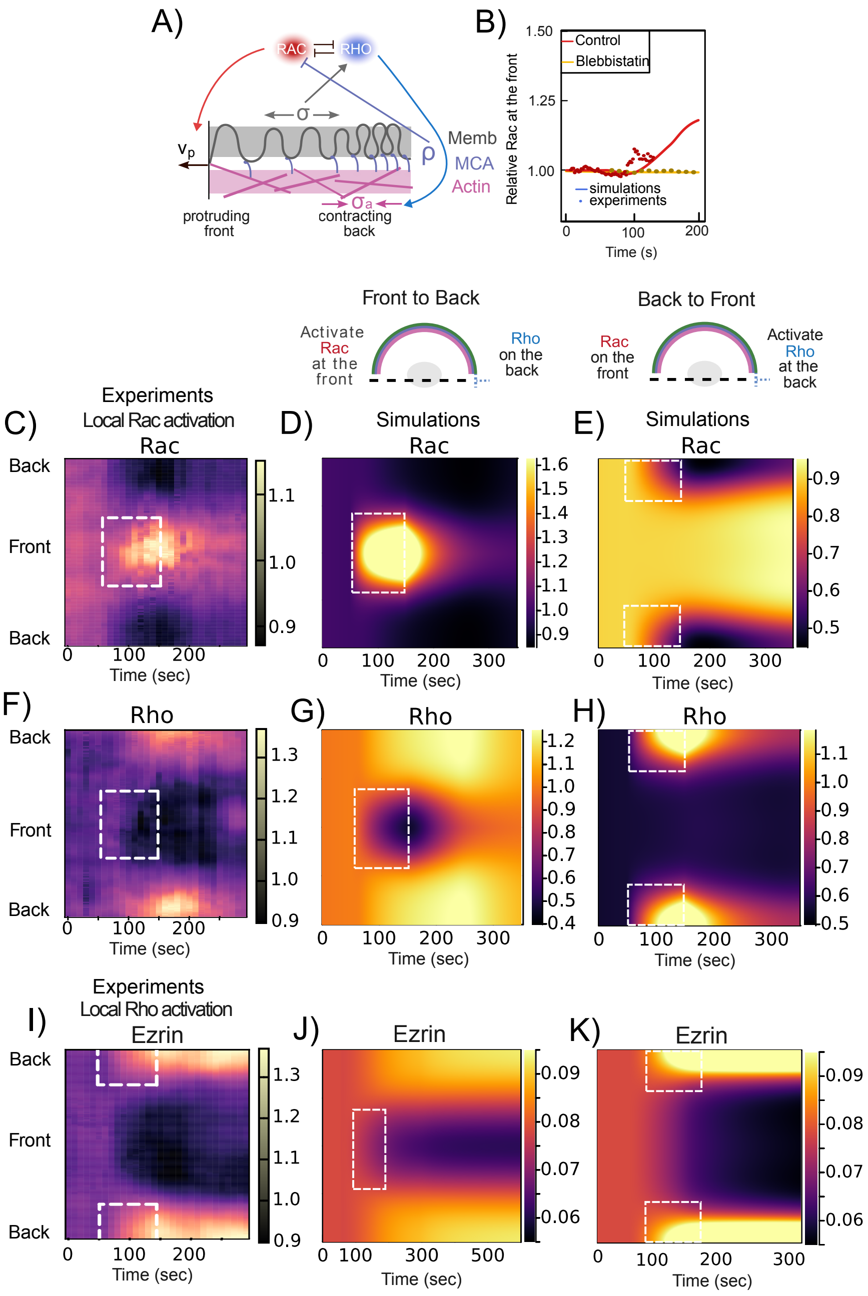
A)
RAC
RHO
σ
Memb
MCA
Actin
ρ
$v_p$
$\sigma_a$
protruding front
contracting back
B)
Relative Rac at the front
Control
Blebbistatin
simulations
experiments
Time (s)
Front to Back
Activate Rac at the front
Rho on the back
Back to Front
Rac on the front
Activate Rho at the back
C)
Experiments
Local Rac activation
Rac
Back
Front
Back
Time (sec)
D)
Simulations
Rac
Time (sec)
E)
Simulations
Rac
Time (sec)
F)
Rho
Back
Front
Back
Time (sec)
G)
Rho
Time (sec)
H)
Rho
Time (sec)
I)
Experiments
Local Rho activation
Ezrin
Back
Front
Back
Time (sec)
J)
Ezrin
Time (sec)
K)
Ezrin
Time (sec)

**<u>Figure S7: Kymographs comparing experimental and simulated data.</u>**

(A) Mechanochemical model of Rac and Rho polarity, combining local mutual inhibition with long-range mechanical feedback. (B) Rho is locally activated on one end (the 'back') of the cell, while EZR (C) and Rac (D) levels are measured at the opposite end (the 'front') of the simulated cell. (Control is shown in red, while cells with impaired contraction (blebbistatin) are shown in yellow. Relative Rac concentration at the front of the cell. The higher discrepancy between model and data here is explained by the fact that Rac activity is dependent on blebbing which varies across cells both in term of timing and numbers of blebs per cells. Here experimental Rac levels are shown only for the first wave of blebbing. Kymographs from experiments and simulations for (D-F) Rac, (G-I) Rho, and (J-L) Ezrin. The first column (D, G, J) corresponds to experiments on both Front to Back and Back to Front. The second column (E, H, K) and third column (F ,I ,L) correspond to simulations results with Front to Back and Back to Front activation respectively. The dotted boxes correspond to the spatiotemporal window of activation of Rac or Rho, for Front to Back and Back to Front respectively.

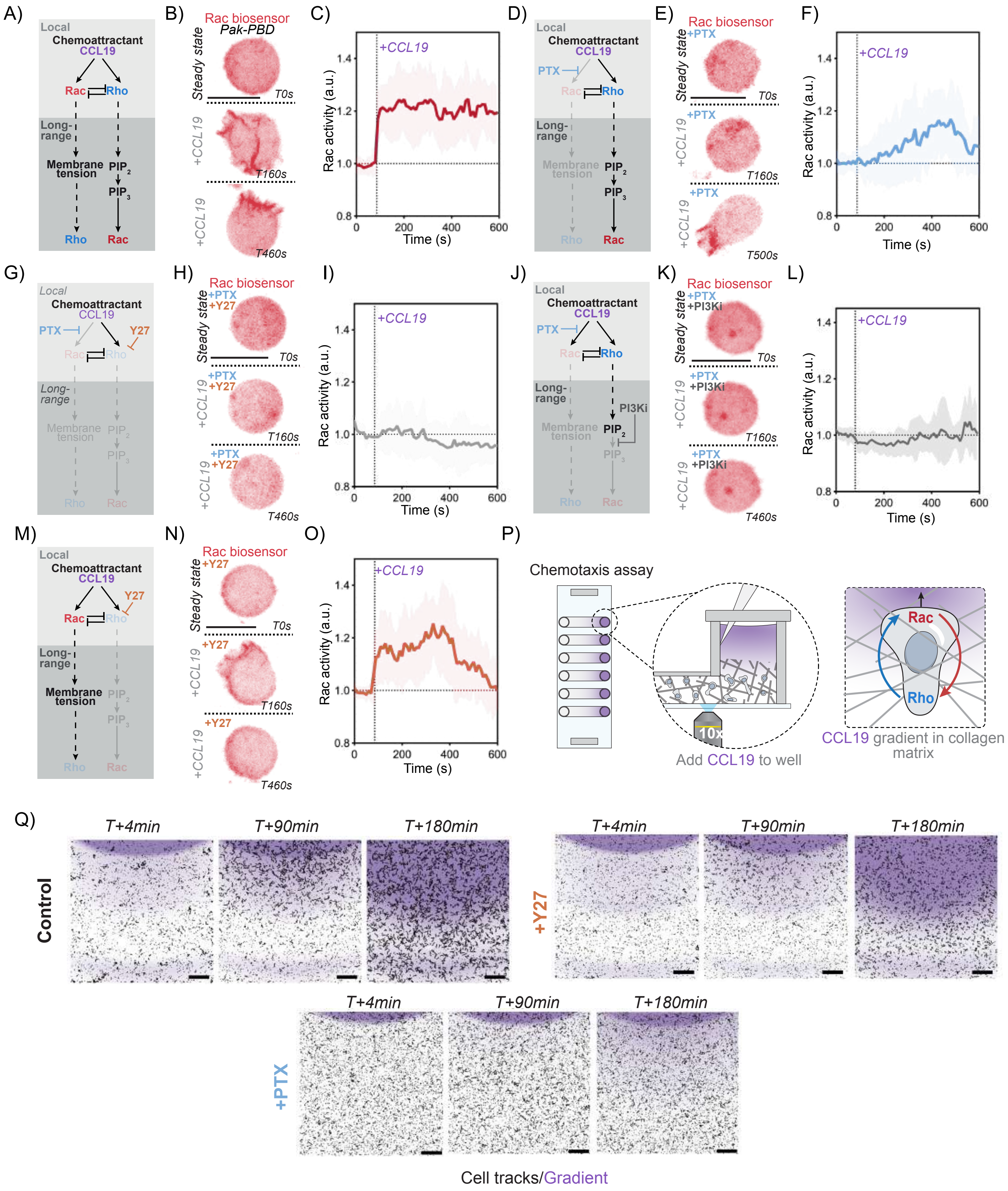

A)
Local
Chemoattractant
CCL19
Rac
Rho
Long-range
Membrane tension
PIP2
PIP3
B)
Rac biosensor
Pak-PBD
Steady state
T0s
+CCL19
T160s
T460s
C)
+CCL19
Rac activity (a.u.)
Time (s)
D)
PTX
E)
+PTX
T500s
F)
G)
Y27
H)
I)
J)
PI3Ki
K)
+PI3Ki
L)
M)
N)
+Y27
O)
P)
Chemotaxis assay
10x
Add CCL19 to well
CCL19 gradient in collagen matrix
Q)
T+4min
T+90min
T+180min
Control
+Y27
+PTX
Cell tracks/Gradient

**Figure S8: Long range mutual activation establishes robust Rho and Rac polarity in primary T cells**

A) Primary human T cells expressing the Rac biosensor (Pak-PBD) to assay cell polarization following acute stimulation with the chemoattractant CCL19. B) CCL19 is known to activate both Rho and Rac. C) Time-lapse confocal images of an unpolarized human primary T cell before and during CCL19 stimulation. Rac activity was monitored via the Rac biosensor Pak-PBD. D) Average time trace of Rac activity at the cell front following addition of CCL19. (mean $\pm$ 95%CI; n>30, N= 3) E) Cells are treated with G$\alpha$I inhibitor PTX to prevent direct Rac activation by CCL19. F) Average time trace of Rac activity at the cell front of cells treated with 1 ug/ml of G$\alpha$I inhibitor following addition of CCL19. (mean $\pm$ 95%CI; n>20, N= 3). G) Cells are treated with a combination of Y27 and PTX to prevent direct Rho and Rac activation by CCL19. H) Average time trace of Rac activity at the cell front of cells treated with 1 ug/ml of G$\alpha$I inhibitor PTX together with 20uM of Y27 following addition of CCL19 (mean $\pm$ 95%CI; n>20, N= 3). I) Cells are treated with a combination of PTX and the PI3K inhibitor Duvelisib to disrupt the long-range Rho-mediated Rac activation described in this study. Average time trace of Rac activity at the cell front of cells treated with 1 ug/ml of G$\alpha$I inhibitor PTX together with 1uM PI3K inhibitor Duvelisib. (mean $\pm$ 95%CI; n>15, N = 2). K) Cells are treated with ROCK inhibitor Y27 to prevent Rho activation. L) Average time trace of Rac activity at the cell front of cells treated with 20uM of Y27 following addition of CCL19 (mean $\pm$ 95%CI; n>20, N= 3).Data for D), F), H) and L) is similar as in **Figure 7D** but separated into different panels for extra clarity. (M) Ex-vivo assay for human primary T cell chemotaxis. Cells are premixed with Bovine Dermal Collagen and placed into linear channels. After collagen sets, media is added to one side of the channel (TCM) and media with human CCL19 (100ng total) and 10ug/mL of Dextran10k-AF647 (Dex647) is added to the other. Imaging takes place right on the edge of the well containing CCL19 so that T cell responses can be recorded as the CCL19 (as read out by Dex647) diffuses into the channel. (N) Representative control, Y27 and PTC experiment at 180s with Dex647 fluorescence and corresponding tracks of T cells; tracks display 3 minutes preceding timepoint listed.

### Supplementary text

Henry De Belly, Andreu Fernández Gallén, Evelyn Strickland, Dorothy C. Estrada, Patrick J. Zager, Janis K Burkhardt, Hervé Turlier, Orion D. Weiner

#### 1 Model hypotheses

Rho GTPases have emerged as key components of the cell polarization machinery. Among them, Rho and Rac are well-known for their antagonistic and inhibitory relationship, often modeled through reaction-diffusion equations [1]. A well-established paradigm for cell polarization is the concept of wave-pinning [2], where bistability of a single RhoGTPase arises from the interplay between a degradation term and a non-linear activation term, combined with overall mass conservation. Recent models have also incorporated the experimentally observed dependence of Rho on membrane tension [3, 4].

In our model, we propose an alternative polarization mechanism to wave-pinning, in which we neglect the shuttling between active and inactive forms of Rho GTPases. Instead, we model the local mutual inhibition between Rho and Rac [5] as a bistable switch, which naturally exhibits both monostable and bistable regimes, depending on the parameter set [6]. The couplings between cortical and membrane mechanics are the essential components that enable cell polarization, and we have designed these based on experimental observations. Similar to previous works [3, 4], and in agreement with our experimental findings, we assume that increased membrane tension promotes RhoA activation.

Our minimal mechanical model integrates the membrane and cortex based on previous work by some of the authors [7]. In this model, membrane tension effectively acts as a tangential elasticity (membrane folding/unfolding), while the cortex is represented as a viscous, contractile layer [8, 9, 10]. Their mechanical interaction is implemented through viscous friction, which slows their relative tangential movement. This leads to effective diffusive propagation of membrane tension along the cell surface [11], which homogenizes in neutrophils after perturbations over timescales longer than several minutes [7].

In our current work, we refine this model by explicitly accounting for the spatiotemporal variations in membrane-cortex attachment (MCA) surface density, particularly of ezrin, which exchanges with its inactive form in the cytoplasm and can be advected by cortical flows. Experimentally, we further observe that protrusions through blebbing are enhanced in regions of the membrane with low MCA and include this key coupling minimally via a switch-like dependence of the protusion velocity with MCA density.

After presenting the local biochemical model for Rho and Rac and analyzing its response to external stimulation, we extend the model to include wave-pinning dynamics and demonstrate the hypersensitivity of this class of models to external forcing. To address the limitations of this approach, we then introduce an alternative mechanochemical model. In this framework, we derive coupled mechanical equations in one dimension and investigate the system's response to external stimulation using the finite-element method.

#### 2 Biochemical model of Rho and Rac

In this section, we first describe the coupled dynamics of Rac ($R$) and Rho ($\rho$), modeled locally via ordinary equations describing their activation modulated by mutual inhibition and their inactivation, in a similar fashion as a toggle-switch [12]. In contrast to wave-pinning models, which account explicitly for the shuttling between an active form of the protein in the membrane and an inactive form in the cytosol, and rely importantly on a limited total amount of RhoGTPase in the cell, we assume here the membrane in contact with a chemiostat of inactive proteins in the cytosol. As such, we assume effectively a constant cytosolic concentration, that is integrated within the activation rates. This simplifying hypothesis is justified by former experimental work, which demonstrated that in neutrophils, a diffusion-based inhibition or local sequestration mechanism was not sufficient to explain polarization [13].

The two ordinary equations read

$$\frac{\partial R}{\partial t} = \alpha_0 \frac{k_R^2}{k_R^2 + \rho^2} + s_R - d_R R, \tag{1}$$

$$\frac{\partial \rho}{\partial t} = \beta_0 \frac{k_\rho^2}{k_\rho^2 + R^2} + s_\rho - d_\rho \rho, \tag{2}$$

where $\alpha_0$ and $\beta_0$ are the basal activation rates of Rac ($R$) and Rho ($\rho$), respectively; $k_R$ and $k_\rho$ set the switch threshold values for $R$ and $\rho$. We introduce source terms for Rac and Rho $s_R$ and $s_\rho$, for any optogenetic or chemotaxis activation of a GPTase.

#### 2.1 Non-dimensionalized equations

We non-dimensionalize previous equations defining dimensionless protein concentrations

$$\bar{\rho} = \rho / k_R \tag{3}$$

$$\bar{R} = R / k_\rho \tag{4}$$

and dimensionless activation rates

$$\bar{\alpha}_0 = \frac{\alpha_0}{k_\rho \, d_R}, \tag{5}$$

$$\bar{\beta}_0 = \frac{\beta_0}{k_R \, d_\rho}. \tag{6}$$

The resulting dimensionless equations for the biochemical equations are

$$\frac{1}{d_R} \frac{\partial \bar{R}}{\partial t} = \bar{\alpha}_0 \frac{1}{1 + \bar{\rho}^2} + \bar{s}_R - \bar{R}, \tag{7}$$

$$\frac{1}{d_\rho} \frac{\partial \bar{\rho}}{\partial t} = \bar{\beta}_0 \frac{1}{1 + \bar{R}^2} + \bar{s}_\rho - \bar{\rho}, \tag{8}$$

#### 2.2 Steady-state solutions

**Nullclines and fixed points**

Steady-state equations ($\partial/\partial t = 0$) define the nullclines

$$0 = \bar{\alpha}_0 \frac{1}{1 + \bar{\rho}^2} - \bar{R} \equiv f(\bar{R}, \bar{\rho}), \tag{9}$$

$$0 = \bar{\beta}_0 \frac{1}{1 + \bar{R}^2} - \bar{\rho} \equiv g(\bar{R}, \bar{\rho}). \tag{10}$$

The intersection of the nullclines $f(\bar{R}, \bar{\rho})$, $g(\bar{R}, \bar{\rho})$ define the possible steady-state solutions as function of the values of activation rates $(\bar{\alpha}_0, \bar{\beta}_0)$. As illustrated on Fig. 1, there are two possible scenarios. Either the nullclines have one intersection - or fixed - point, which is always a stable solution, and corresponds either to a (high Rac/low Rho), or (low Rac/high Rho) situation; or the curves intersect on three fixed points, two of which are stable solutions while the intermediate one is unstable. In the former case, the system is said monostable, while in the latter, the system displays a bistable behavior where both (high Rac/low Rho) and (low Rac/high Rho) coexist as possible stable solutions.

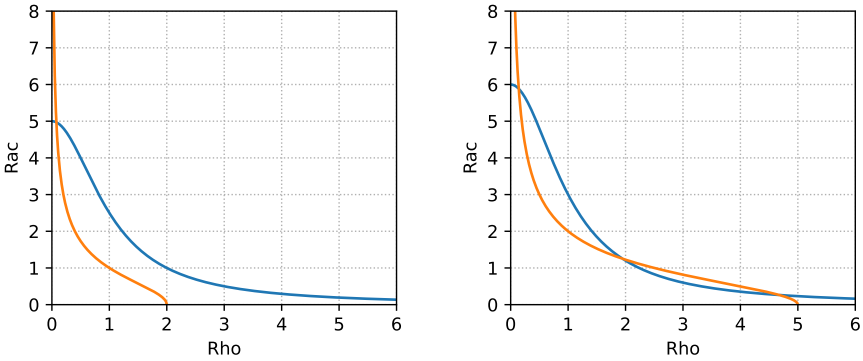


Figure 1: Plot of eqs (9) and (10) for the values $\bar{\alpha}_0 = 5$ $\bar{\beta}_0 = 2$ and $\bar{\alpha}_0 = 6$ $\bar{\beta}_0 = 5$ respectively. There is a steady state solution for $R$ and $\rho$ where this two lines cross.

##### Phase diagram

We can compute the regions of parameters $(\bar{\alpha}_0, \bar{\beta}_0)$ of monostability and bistability by looking for the roots $(\bar{R}^*, \bar{\rho}^*)$ of the steady-state equations $(f(\bar{R}, \bar{\rho}) = 0,\ g(\bar{R}, \bar{\rho}) = 0)$, while ensuring that these roots are stable fixed points $(\partial_{\bar{R}} f(\bar{R}, \bar{\rho})|_{\bar{R}^*, \bar{\rho}^*} < 0\ \partial_b g(\bar{R}, \bar{\rho})|_{\bar{R}^*, \bar{\rho}^*} < 0)$.

The transition lines between monostable and bistable stability regions are obtained when

$$f(\bar{R}, \bar{\rho}) = 0, \quad g(\bar{R}, \bar{\rho}) = 0, \quad \partial_{\bar{R}} f(\bar{R}, \bar{\rho}) = 0, \quad \partial_{\bar{\rho}} g(\bar{R}, \bar{\rho}) = 0, \tag{11}$$

which leads to the following polynomial equations for $(\bar{R}, \bar{\rho})$

$$0 = -\bar{\alpha}_0 + \bar{R}\left(1 + \bar{\beta}_0^2\right) - 2\,\bar{R}^2\bar{\alpha}_0 + 2\,\bar{R}^3 - \bar{R}^4\bar{\alpha}_0 + \bar{R}^5 \tag{12}$$

$$0 = -\bar{\beta}_0 + \bar{\rho}\left(1 + \bar{\alpha}_0^2\right) - 2\,\bar{\rho}^2\,\bar{\beta}_0 + 2\,\bar{\rho}^3 - \bar{\rho}^4\,\bar{\beta}_0 + \bar{\rho}^5 \tag{13}$$

The phase diagram is plotted on Fig. 2 as function of $\bar{\alpha}_0$ and $\bar{\beta}_0$.

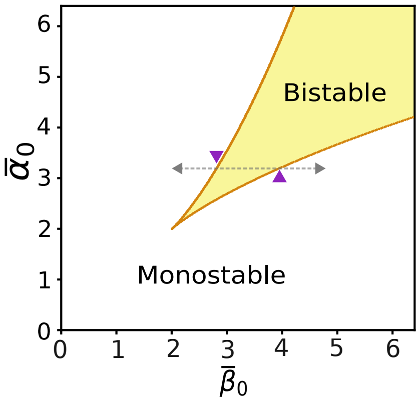


Figure 2: Phase diagram of mono and bi-stability regions of Rho and Rac, as function of their normalized activation rates $\bar{\beta}_0$ and $\bar{\alpha}_0$. The transition curves are in orange. Plotted in grey arrow the hysteresis represented in Figure 3 for the line $\bar{\alpha} = 3.2$ and represented and with purple triangles each transition point for the hysteresis.

The transition from low to high Rho/Rac and conversely exhibits a hysteresis, that we illustrate on Fig. 3. At a given value $\bar{\alpha}_0 = 3.2$, if one increases $\bar{\beta}_0$ from a low value $\sim 0$, the system will switch from (high Rac, low Rho) to (low Rac, high Rho) only when $\bar{\beta}_0$ reaches the second transition curve at $\bar{\beta}_0 \simeq 3.92$. Traversing the parameter space in the reverse direction, starting from $\bar{\beta}_0 \sim 6$ and decreasing its value the transition from (low Rac, high Rho) will happen this time when goes below $\bar{\beta}_0 \simeq 2.82$

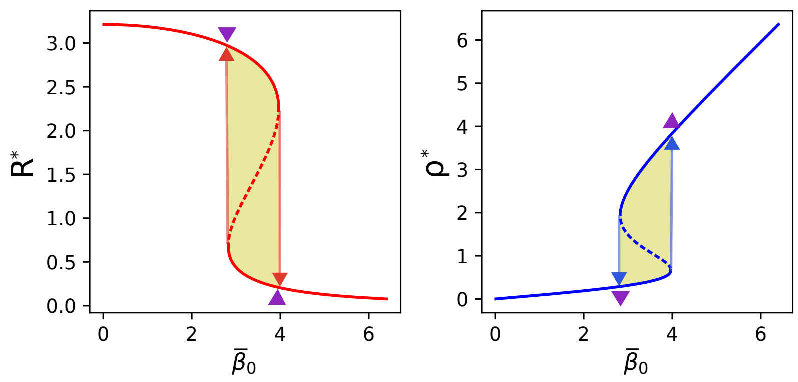


Figure 3: Steady-state solutions of Rac $\bar{R}^*$ and Rho $\bar{\rho}^*$ as function of $\bar{\beta}_0$ ($\bar{\alpha}_0 = 3.2$ being constant). The solid lines represent stable solutions while the dotted line represents unstable solutions. The hysteresis is materialized through arrows, which delimits the bistability region in yellow $\bar{\beta}_0 \in [2.82, 3.92]$. The purple triangles pointing up and down correspond to the transition line points depicted on Fig. 2.

#### 2.3 Biochemical model with external stimulation: hysteresis and sensitivity to initial conditions

To model the optogenetic stimulation (or inhibition) of the two GTPase activity, we add source/stimulation (i.e. forcing) terms $\bar{s}_R$, $\bar{s}_\rho$ to our biochemical model:

$$\frac{1}{d_{\bar{R}}}\frac{\partial \bar{R}}{\partial t} = \bar{\alpha}_0 \frac{1}{1+\bar{\rho}^2} + \bar{s}_R - \bar{R}, \tag{14}$$

$$\frac{1}{d_{\bar{\rho}}}\frac{\partial \bar{\rho}}{\partial t} = \bar{\beta}_0 \frac{1}{1+\bar{R}^2} + \bar{s}_\rho - \bar{\rho}. \tag{15}$$

Starting from a given stationary stable state in the bistable region, the external (possibly transient) activation (or inhibition) of one GTPase can make the stationary solution switch to the other stable state. This switch depends on the initial solution and on the type and amplitude of the forcing, which is a manifestation of the hysteresis, as illustrated on Fig. 4.

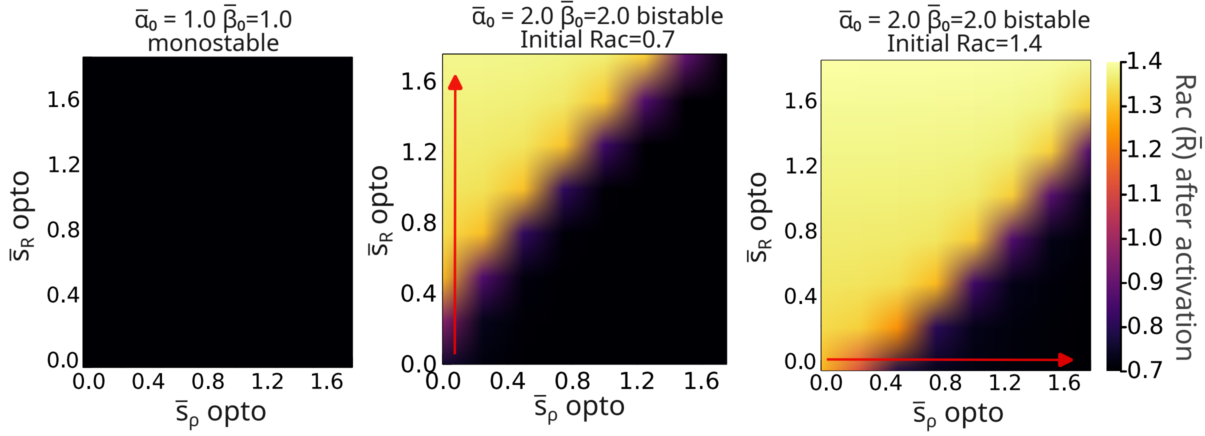


Figure 4: Heatmap of adimensional Rac concentration $\bar{R}$ at the end of the simulation as a function of Rac and Rho stimulation intensities $\bar{s}_R$ and $\bar{s}_\rho$ in a point-like system. In the bistable regime, the outcome depends on the initial value of $\bar{R}$. During the simulation, localized Rac and Rho stimuli are applied at the same spatial point for 100 seconds. After stimulus removal, the system relaxes, and the final value of $\bar{R}$ is plotted. Arrows indicate the directions of the most effective stimuli for switching Rac from the zero state $\bar{s}_R = \bar{s}_\rho = 0$.

The figure 4 illustrates how Rac behaves in response to an external stimulation under distinct basal production rates:

- In the monostable parameter regime (left panel), Rac always converges to the same steady-state value, regardless of the external activation of Rac or Rho (i.e., regardless of the values of $\bar{s}_R$ or $\bar{s}_\rho$).
- In the bistable parameter regime (center and right panels), Rac can switch from one stable steady-state value $R^*$ to the other through transient activation of either Rac or Rho (via $\bar{s}_R$ or $\bar{s}_\rho$ respectively).

In the bistable regime, if the system starts at the lower steady-state value of Rac, applying a sufficiently strong Rac activation ($\bar{s}_R$) can induce a switch to the higher Rac state (center panel, Fig. 4). However, if the system is initially in the higher Rac state, further Rac activation will not lead to a switch. In this case, transitioning back to the lower state requires Rho activation ($\bar{s}_\rho$), which inhibits Rac and allows the system to fall to the lower stable solution (third panel, Fig. 4).

In summary, in a purely biochemical model, the ability to switch between stable states depends strongly on the initial conditions and which GTPase is activated. Activating only one GTPase may not always be sufficient to induce a state transition. This poses a limitation for robust modeling of cell polarization. However, this issue is resolved in the mechanochemical spatially extended model, where mechanical feedback enables consistent and sustained polarization independently of the initial state.

#### 2.4 Spatially extended biochemical model

The previous model describes GTPase dynamics at a single point in space, without accounting for their spatial distribution within the cell. To incorporate spatial effects, a mechanism of transport must be introduced. As a first step, we consider the simplest form of transport: the diffusion of GTPases along the cell membrane.

$$\frac{1}{d_{\bar{R}}}\frac{\partial \bar{R}}{\partial t} = \bar{\alpha}_0 \frac{1}{1+\bar{\rho}^2} + \bar{s}_R - \bar{R} + \bar{D}_R \nabla^2 \bar{R}, \tag{16}$$

$$\frac{1}{d_{\bar{\rho}}}\frac{\partial \bar{\rho}}{\partial t} = \bar{\beta}_0 \frac{1}{1+\bar{R}^2} + \bar{s}_\rho - \bar{\rho} + \bar{D}_\rho \nabla^2 \bar{\rho}. \tag{17}$$

Adding diffusion terms to the biochemical system in the bistable regime, and initializing it from a polarized state (with front and back corresponding to steady-state solutions), results in a polarization front that gradually dissipates over time (see Fig. 5). This simple spatially extended biochemical model is therefore insufficient to explain sustained cell polarization, as it fails to maintain a stable polarized state.

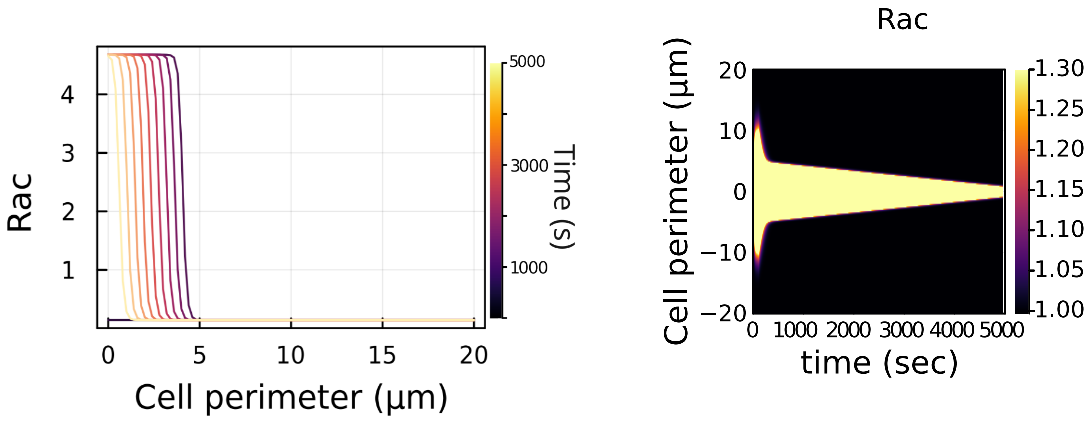


Figure 5: Absence of steady-state solutions in the presence of diffusion. Left: Successive spatial profiles of membrane-bound Rac concentration $\bar{R}$ over time, showing the decay of polarization. Right: Kymograph of $\bar{R}$, with time on the x-axis and spatial position on the y-axis, illustrating the progressive flattening of the Rac concentration profile.

To overcome this limitation, a class of models known as wave-pinning models has been proposed. These models introduce the key assumption of mass conservation between the active and inactive forms of GTPases, which enables the stabilization of the polarization front. We introduce this framework in the following section.

### 3 Wave pinning model

We generalize below our spatially extended biochemical model to explicitly account for the reversible switching between active and inactive states of GTPases, while assuming a fixed total amount for each protein. This approach follows the Wave-Pinning (WP) framework introduced in [2].

In this context, bistability for a single diffusive Rho GTPase emerges from the interplay between a nonlinear activation term and the constraint of mass conservation between the active and inactive forms. The governing equations for this system are:

$$\frac{1}{d_R}\frac{\partial \bar{R}}{\partial t} = \bar{\alpha}_0\,\frac{1}{1+\bar{\rho}^2}\bar{R}_i + \bar{s}_R - \bar{R} + D_R\nabla^2\bar{R} \tag{18a}$$

$$\frac{1}{d_\rho}\frac{\partial \bar{\rho}}{\partial t} = \bar{\beta}_0\,\frac{1}{1+\bar{R}^2}\bar{\rho}_i + \bar{s}_\rho - \bar{\rho} + D_\rho\nabla^2\bar{\rho}. \tag{18b}$$

$$\frac{1}{d_R}\frac{\partial \bar{R}_i(\xi,t)}{\partial t} = -\bar{\alpha}_0\,\frac{1}{1+\bar{\rho}^2}\bar{R}_i - \bar{s}_R\ + \bar{R} + D_{Ri}\nabla^2\bar{R} \tag{18c}$$

$$\frac{1}{d_\rho}\frac{\partial \bar{\rho}_i(\xi,t)}{\partial t} = -\bar{\beta}_0\,\frac{1}{1+\bar{R}^2}\bar{\rho}_i - \bar{s}_\rho\ + \bar{\rho} + D_{\rho i}\nabla^2\bar{\rho}. \tag{18d}$$

where $R_i$ and $\rho_i$ denote the cytoplasmic concentrations of inactive Rac and Rho GTPases at the membrane. Their diffusion coefficients, $D_{\rho i}$ and $D_{Ri}$, are assumed to be much larger than those of their active membrane-bound counterparts, reflecting the faster diffusion of GTPases in the cytoplasm.

To account for mass conservation, we define the total number of moles of Rac and Rho respectively (active + inactive) as follows

$$\int \Big(R(\xi) + R_i(\xi)\Big)d\xi = N_R, \tag{19}$$

and

$$\int \Big(\rho(\xi) + \rho_i(\xi)\Big)d\xi = N_\rho, \tag{20}$$

Based on the assumption that inactive GTPases diffuse rapidly, we approximate their spatial profiles as uniform. Specifically, we write:

$$R_i(\xi) \simeq \langle R_i\rangle, \tag{21}$$

$$\rho_i(\xi) \simeq \langle \rho_i\rangle, \tag{22}$$

where the angle brackets $\langle\cdot\rangle$ denote the spatial average over the domain. From this approximation, we deduce the following mass conservation relation:

$$\langle R_i\rangle \cdot L = N_R - \int R(\xi)\,d\xi, \tag{23}$$

where L is the total membrane length. An analogous relation holds for the Rho GTPase system:

$$\langle \rho_i\rangle \cdot L = N_\rho - \int \rho(\xi)\,d\xi. \tag{24}$$

This leads to an updated set of equations that incorporate mass conservation explicitly:

$$\frac{1}{d_R}\frac{\partial \bar{R}}{\partial t} = \bar{\alpha}_0\,\frac{1}{1+\bar{\rho}^2}\langle\bar{R}_i\rangle\ + \bar{s}_R - \bar{R} + D_R\nabla^2\bar{R}, \tag{25a}$$

$$\frac{1}{d_\rho}\frac{\partial \bar{\rho}}{\partial t} = \bar{\beta}_0\,\frac{1}{1+\bar{R}^2}\langle\bar{\rho}_i\rangle + \bar{s}_\rho - \bar{\rho} + D_\rho\nabla^2\bar{\rho}, \tag{25b}$$

$$\langle\bar{R}_i\rangle\ = \frac{1}{L}\Big(\bar{N}_R - \int \bar{R}(\xi)\,d\xi\Big), \tag{25c}$$

$$\langle\bar{\rho}_i\rangle = \frac{1}{L}\Big(\bar{N}_\rho - \int \bar{\rho}(\xi)\,d\xi\Big). \tag{25d}$$

##### 3.1 Implementation of the wave-pinning model

The wave-pinning (WP) model demonstrates bistability and the capacity to polarize, as illustrated in Fig. 6 for a specific set of parameters.

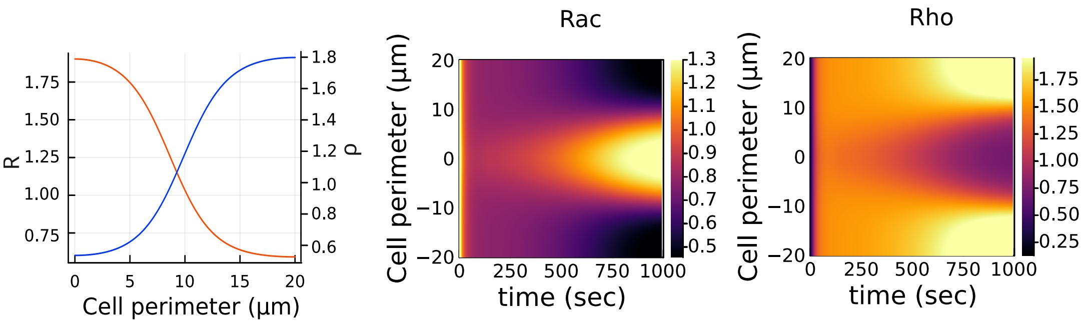


Figure 6: Wave pinning simulations using $\bar{N}_R = 2$ and $\bar{N}_\rho = 2$ and $\bar{\alpha} = \bar{\beta} = 3.0$. Here we have the final profiles of Rac and Rho ($R$ and $\rho$) as well as their kymographs over the cell perimeter and time.

However, the bistable region in the extended system becomes highly dependent on the total amount of Rac and Rho, denoted by $\bar{N}R$ and $\bar{N}\rho$, in addition to their respective production rates, $\bar{\alpha}_0$ and $\bar{\beta}_0$, as shown in Figure 7. For the system to polarize, the total number of moles of each GTPase must be sufficiently large; moreover, the bistable region expands with increasing total mass. Interestingly, each GTPase exhibits a relatively independent bistable parameter region, determined primarily by its own production rate and total mass. This occurs largely independently of the mutual inhibition between Rho and Rac observed in the point-like system, which effectively becomes negligible in the spatially extended context.

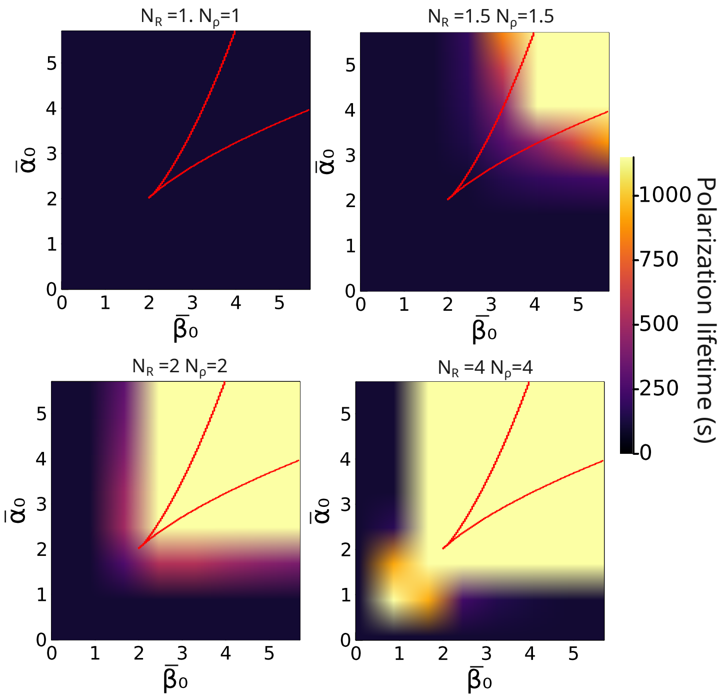


Figure 7: Heatmap of polarization lifetime in the wave-pinning model as a function of the dimensionless production rates of Rac and Rho, $\bar{\alpha}_0$ and $\bar{\beta}0$, for four different values of total Rac and Rho levels, $N_R$ and $N\rho$.

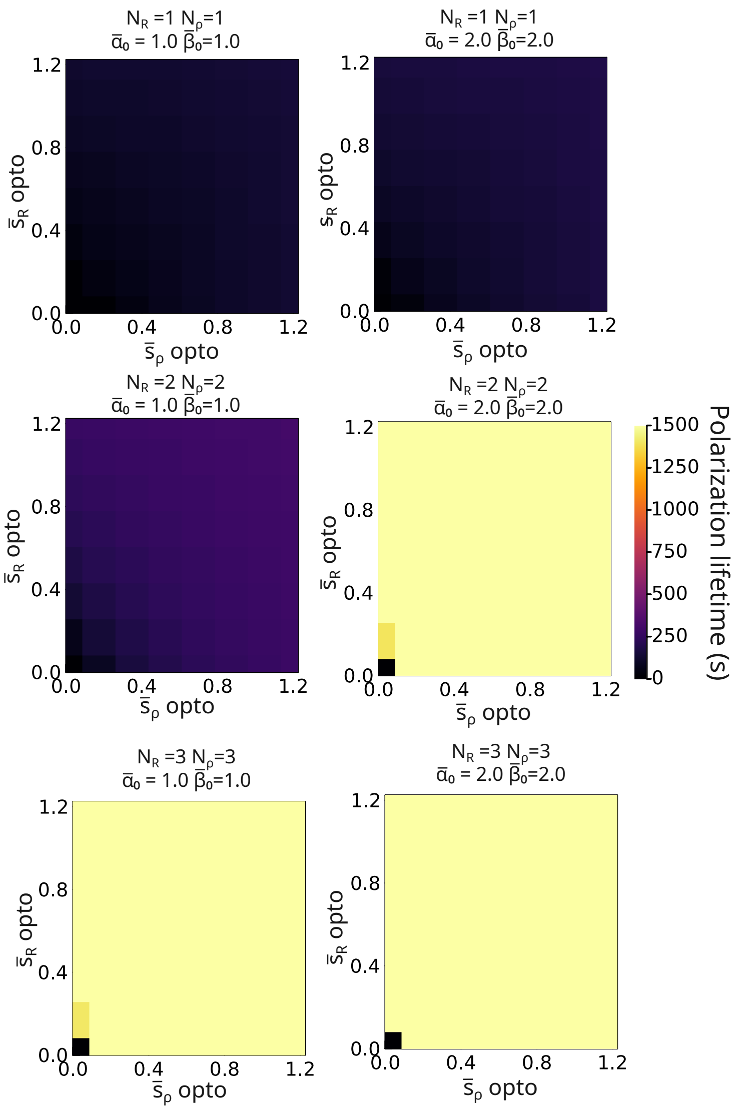


Figure 8: Heatmap of polarization lifetime in the wave-pinning model as a function of the induced source terms for Rac and Rho, $\bar{s}R$ and $\bar{s}\rho$, for three different values of total Rac and Rho levels ($N_R$ and $N_\rho$). The Rac source is applied at the front of the cell, while the Rho source is applied at the back. Source activation lasts for 100 seconds, after which the polarization lifetime is measured.

#### 3.2 Polarization lifetime under external stimulation for the wave-pinning model

The source terms $\bar{s}R$ and $\bar{s}\rho$ can be used to simulate optogenetic or chemotactic activation of Rac and Rho within a cell. In Fig. 8, we examine the persistence of polarization following optogenetic activation under various conditions, while keeping the production rates of Rac and Rho ($\bar{\alpha}_0$ and $\bar{\beta}_0$) fixed.

The results shown in Figure 8 highlight key features of Rac-Rho polarization under the WP model. First, polarization is highly sensitive to the total amounts of Rac and Rho ($N_R$ and $N_\rho$), with polarization failing to occur below a critical threshold. Second, when the WP model allows for polarization under external stimulation, it displays a binary response to external stimuli—either minimal response or full polarization—with little evidence of intermediate states. Notably, under polarizable conditions, even weak activation is sufficient to trigger a polarized state, that would lead to a generic hypersensitivity of the cell. This extreme sensitivity may present a limitation. In biological systems, one might expect that polarization under permissive conditions would require a stimulus of sufficient magnitude to prevent spontaneous activation by stochastic fluctuations.

##### 3.3 Limitations of wave pinning models for GTPase polarization

The proposed system achieves polarization through a diffusion-driven mechanism coupled with GTPase mass conservation. However, experimental evidence suggests that membrane diffusion alone may be insufficient to drive polarization in cells [13]. Therefore, while the VP model demonstrates the capacity for polarization, this mechanism may not be physiologically sufficient on its own.

Moreover, our experimental observations reveal that Rac activation is associated with membrane blebbing, whereas Rho activation correlates with increased membrane tension. These insights motivate the development of an integrated model that incorporates mechanical cues alongside the biochemical dynamics of GTPases.

A further limitation of the WP model is its excessive sensitivity to external stimuli: even minor inputs can readily induce polarization. Given the inherently noisy nature of biological systems, such hypersensitivity could be unrealistic: robust cellular polarization should require a threshold level of stimulation to prevent spurious activation by fluctuations.

#### 4 Mechanical model

##### 4.1 Hypotheses

To describe the mechanics of the membrane - cortex interaction, we follow a previous composite model [7] that we expand to explicitly account for the inhomogeneous membrane-cortex attachment density (MCA) revealed experimentally (Supp Fig. 5A-D). The membrane mechanics is approximated to a linear elastic response at the macroscopic level, resisting effectively stretching or compressive stress through a membrane tension $\sigma$, which may hence be positive or negative. This effective tension is the result of the highly folded structure of the membrane, that results physically from a competition between thermal and active fluctuations, bending rigidity and compressive stress induced by cortical contraction. The cortex is modeled classically as a viscous and contractile shell-like material on the timescales of our experiments $t \simeq 100s$, described by a velocity $v$ and an active contractile stress $\sigma_a$. The membrane and cortex interact through MCA proteins, such as ezrin, that are firmly attached to the cortex in their active state but can flow within the membrane plane, that is tangentiallly fluid. The relative movement between the membrane and cortex results therefore in a friction term that is proportional to the relative velocity between the two surfaces and to to the local surface density of MCA proteins. We further account for the binding kinetics of MCA proteins from the membrane to the cortex and therefore distinguish the density of bound proteins $\rho_b$ from unbound density $\rho_u$. The binding of ERM MCA proteins such as ezrin is regulated by their phosphorylation state, it binds to the cortex in the active form and unbinds when it is dephosphorylated [14]. The model components are summarized in the sketch on Fig. 9.

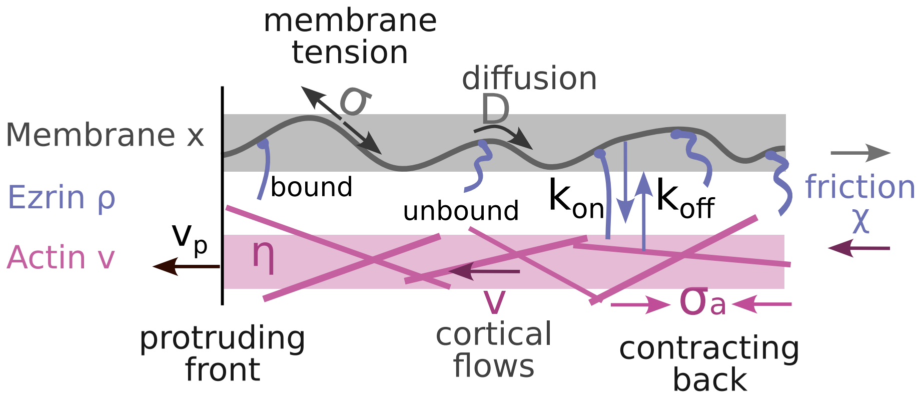


Figure 9: Representation of the mechanical model.

##### 4.2 General equations

Let $x$ denote the membrane strain from its relaxed - or reference - state, and $\dot{x} = (\partial x)/(\partial t)$ its Eulerian velocity. We introduce $\xi\,[N.s.m^{-1}]$ the friction per bound ezrin protein in the membrane and $\sigma_0[N.m^{-1}]$

the two-dimensional membrane elastic modulus associated to its folding/unfolding, and which is homogeneous to a tension. The friction density per unit surface is given by $\rho_b \xi \, [N.s.m^{-3}]$, where $\rho_b \, [m^{-2}]$ is the surface density of bound ezrin proteins. Effective tangential force balance in the membrane reads

$$\sigma_0 \nabla^2 x - \chi \rho_b \Big( \dot{x} - v \Big) = 0. \tag{26}$$

Let $v$ denote the local tangential velocity of the cortex. We introduce $\eta \, [N.s.m^{-1}]$ the two-dimensional effective viscosity of the cortex and $\sigma_a \, [N.m^{-1}]$ the two-dimensional active tension in the cortex, regulated by the local actomyosin contractility. Tangential force balance in the cortex reads

$$\eta \nabla^2 v + \nabla \sigma_a - \chi \rho_b \Big( v - \dot{x} \Big) = 0. \tag{27}$$

We note that the instantaneous local membrane tension may be defined from the local stretch as

$$\sigma = \sigma_0 \nabla x \tag{28}$$

We furthermore introduce the conservation equations for bound and unbound MCA proteins $\rho_b$ and $\rho_u$ at the surface as

$$\frac{\partial \rho_b}{\partial t} + \nabla (v \rho_b) = k_{\text{on}} \rho_u - k_{\text{off}} \rho_b - \lambda_b \rho_b^3, \tag{29a}$$

$$\frac{\partial \rho_u}{\partial t} + \nabla \big( \dot{x} \rho_u \big) = D \nabla^2 \rho_u - k_{\text{on}} \rho_u + k_{\text{off}} \rho_b, \tag{29b}$$

where we introduced the binding and unbinding rates $k_{\text{on}} \, [s^{-1}]$ and $k_{\text{off}} \, [s^{-1}]$, a two-dimensional diffusion coefficient $D \, [m.s^{-1}]$ for the unbound ezrin within the membrane plane and a non-linear saturation term for the bound protein density $\lambda_b$.

#### 4.3 One-dimensional formulation

To simplify the description of the cortex-membrane interaction and exploiting the axisymmetry of the problem, we approximate in the following the system as a one-dimensional string of length $L$. We don't expect any of our results to be qualitatively different than with a more realistic 2D or 3D model of the surface. The composite surface deformation is described in one-dimension with respect to a spatial coordinate $\xi$, such that previous momentum and mass conservation equations read

$$\sigma_0 \frac{\partial^2 x}{\partial \xi^2} - \chi \rho_b (\dot{x} - v) = 0, \tag{30a}$$

$$\eta \frac{\partial^2 v}{\partial \xi^2} + \nabla \sigma_a - \chi \rho_b \Big( v - \dot{x} \Big) = 0, \tag{30b}$$

$$\frac{\partial \rho_b}{\partial t} + \frac{\partial (v \rho_b)}{\partial \xi} = k_{\text{on}} \rho_u - k_{\text{off}} \rho_b - \lambda_b \rho_b^3, \tag{30c}$$

$$\frac{\partial \rho_u}{\partial t} + \frac{\partial (\dot{x} \rho_u)}{\partial \xi} = D \frac{\partial^2 \rho_u}{\partial \xi^2} - k_{\text{on}} \rho_u + k_{\text{off}} \rho_b, \tag{30d}$$

These equations need to be complemented by appropriate boundary and intial conditions. We assume that the protrusive front can be modeled through a Dirichlet boundary conditions as a protrusive velocity $v_p$ for the cortex and a corresponding deformation due to protrusions for the membrane at $\xi = 0$. Since our one-dimensional system shall be seen as a representation of an axisymmetric mmembrane, "unwrapped" into a one-dimensional string, we furthermore impose zero Dirichlet boundary conditions for both the membrane strain and cortex velocity by symmetry at the opposite side of the cell (back) $\xi = L$:

$$v(\xi = 0) = v_p \, , \quad \text{and} \quad v(\xi = L) = 0 \tag{31}$$

$$x(\xi = 0) = \int_0^t v_p(t) \, dt. \, , \quad \text{and} \quad x(\xi = L) = 0 \tag{32}$$

One drawback of the simplified one-dimensional formulation is that the membrane may not relax even when the velocity reaches zero. To address this issue, we introduced an additional slow linear relaxation over time for $x(\xi = 0)$, accounting not only for protrusion retraction but also for other membrane relaxation mechanisms, such as membrane addition via endocytosis or exocytosis.

For ezrin surface density, we assume zero flux boundary conditions at membrane edges by symmetry again, which translate into zero Neumann boundary conditions at $\xi = 0$ and $\xi = L$

$$\frac{\partial \rho_b}{\partial \xi}\Big|_{0,L} = 0 \quad \text{and} \quad \frac{\partial \rho_u}{\partial \xi}\Big|_{0,L} = 0. \tag{33}$$

The initial values of bound and unbound densities are homogeneous and set such that their sum equals a conserved total number of ezrin proteins, defined as $M_\rho = \int (\rho_b + \rho_u) d\chi$, which allows us to define a characteristic mean ezrin density in one dimension

$$\rho_0 \equiv \frac{M_\rho}{L} \tag{34}$$

Unless specified, all other variables are set initially to their homogeneous basal or stationary values.

Although the parameters in the one-dimensional formulation of the model have different dimensions than their two-dimensional counterparts, the dimensionless parameters we will define further remain unchanged, and their numerical values can therefore be directly inferred from experimental measurements.

### 5 Mechanochemical model

#### 5.1 Mechanochemical couplings

The mechanochemical model combines the local biochemical model for Rho and Rac described in Section 2 and the composite membrane-cortex mechanics described in Section 4. The control of Rho and Rac on cortical contractility and protrusion activity respectively are well described in the literature [15]. The feedbacks of mechanics on biochemical regulation are in contrary formulated based on experimental results from this manuscript.

**Cortical tension**
The dependence of the contractile tension on Rho concentration in the membrane is assumed to be linear:

$$\sigma_a = \sigma_a^0 \, \frac{b}{k_R}, \tag{35}$$

where $\sigma_a^0$ is a basal contractile tension in the cortex.

**Protrusive velocity**
For the protrusive activity, we assume that the protrusion velocity is activates with Rac and introduce a switch mechanism where protrusion occurs above a Rac concentration threshold $R_{\text{th}}$ , while also accounting for the resistance exerted by membrane tension $\sigma$ [16]

$$v_p \simeq \frac{v_0}{(1 + \sigma^2/\sigma_0^2)} \text{th}\big(R - R_{\text{th}}\big), \tag{36}$$

$$\text{th}\big(R - R_{\text{th}}\big) = \frac{1}{2}\Big[\tanh\Big(\frac{R - R_{\text{th}}}{k_R}\Big) + 1\Big]. \tag{37}$$

**Rac activation**
Based on experimental results on Fig. 4, we furthermore assume that the Rac activation rate depends on the bound ezrin concentration $\rho_b$ in a switch like manner, where it is increased below a threshold value $\rho_{\text{th}}$. We do not model explicitly stochastic blebbing and the competition for Pip2 for simplicity. The spatially-resolved equation for Rac dynamics is modified accordingly as follows

$$\frac{\partial R}{\partial t} = \Big(\alpha_0 + \alpha \, \text{th}(\rho_b - \rho_{\text{th}})\Big) \frac{k_R^2}{k_R^2 + \rho^2} - d_R R + D_R \nabla^2 R, \tag{38}$$

$$\text{where } \text{th}(\rho_b - \rho_{\text{th}}) = \frac{1}{2}\Big[1 - \tanh\Big(\frac{\rho_b - \rho_{\text{th}}}{\rho_0}\Big)\Big], \tag{39}$$

where we introduced a membrane diffusion coefficient $D_R$ for Rac, a basal value $\rho_0$ for the ezrin surface density that will be defined later, and an additional activation rate $\alpha$ for the dependence on ezrin.

**Rho activation**

Based on experimental results on Fig. 2E-L, we finally assume that the Rho activation rate depends on the membrane tension $\sigma$ in a switch like manner, where it is increased above a threshold value $\sigma_{\rm th}$. The spatially-resolved equation for Rho dynamics is modified accordingly as follows

$$\frac{\partial \rho}{\partial t} = \Big(\beta_0 + \beta\,\mathrm{th}(\sigma - \sigma_{th})\Big)\frac{k_\rho^2}{k_\rho^2 + R^2} - d_\rho \rho + D_\rho \nabla^2 \rho \tag{40}$$

$$\text{where } \mathrm{th}(\sigma - \sigma_{\rm th}) = \frac{1}{2}\Big[\tanh\Big(\frac{\sigma - \sigma_{\rm th}}{\sigma_0}\Big) + 1\Big], \tag{41}$$

where we introduced a membrane diffusion coefficient $D_\rho$ for Rho, and an additional activation rate $\beta$ for the dependence on membrane tension.

#### 5.2 Coupled dynamic equations

The full mechanochemical model has six dynamical variables $R$, $\rho$, $v$, $x$, $\rho_b$ and $\rho_u$, the dynamics of which is governed by six partial differential equations

$$\frac{\partial R}{\partial t} = \Big(\alpha_0 + \alpha\,\mathrm{th}(\rho_b - \rho_{\rm th})\Big)\frac{k_R^2}{k_R^2 + \rho^2} - d_R R + D_R \nabla^2 R, \tag{42a}$$

$$\frac{\partial \rho}{\partial t} = \Big(\beta_0 + \beta\,\mathrm{th}(\sigma - \sigma_{th})\Big)\frac{k_\rho^2}{k_\rho^2 + R^2} - d_\rho \rho + D_\rho \nabla^2 \rho, \tag{42b}$$

$$\sigma_0 \nabla^2 x - \chi\,\rho_b\Big(\dot{x} - v\Big) = 0, \tag{42c}$$

$$\eta \nabla^2 v + \sigma_a^0 \nabla\left(\frac{b}{k_R}\right) - \chi\,\rho_b\Big(v - \dot{x}\Big) = 0, \tag{42d}$$

$$\frac{\partial \rho_b}{\partial t} + \nabla(v \cdot \rho_b) = k_{\rm on}\rho_u - k_{\rm off}\rho_b - \lambda_b \rho_b^3, \tag{42e}$$

$$\frac{\partial \rho_u}{\partial t} + \nabla\big(\dot{x} \cdot \rho_u\big) = -k_{\rm on}\rho_u + k_{\rm off}\rho_b + D\nabla^2 \rho_u, \tag{42f}$$

To these equations, one has to add the boundary conditions, which have been made explicit in one dimension in (32), (31), (33). Additional zero-flux boundary conditions for Rac and Rho surface density $R$ and $\rho$ can be expressed identically to those for $\rho_b$ and $\rho_u$ (33) as Neumann boundary conditions. Finally, the explicit coupling of the protrusion velocity with other variables is defined in (36).

#### 5.3 Non-dimensionalization

We non-dimensionalize the previous equations as follows:

We set a basal contractile tension in the cortex $\sigma_a^0$, which allows us to non-dimensionalize the membrane tension and to define a characteristic timescale associated to the active-viscous cortex relaxation that will serve to non-dimensionalize all other times

$$\bar{\sigma}_0 = \frac{\sigma_0}{\sigma_a^0}, \quad \tau_a \equiv \frac{\eta}{\sigma_a^0}, \quad \bar{t} = \frac{t}{\tau_a}. \tag{43}$$

We define a hydrodynamic length $\lambda$ measuring the spatial extent of cortical viscous flows slowed down by friction and we non-dimensionalize all lengths by a characteristic size $L$ of the cell

$$\lambda \equiv \sqrt{\frac{\eta}{\chi\,\rho_0}}, \quad \bar{\lambda} = \frac{\lambda}{L}, \quad \bar{\xi} = \frac{\xi}{L}, \quad \bar{x} = \frac{x}{L}, \quad \bar{v} = v\frac{\tau_a}{L}. \tag{44}$$

Ezrin surface densities are non-dimensionalized using the mean density $\rho_0$ defined in (34)

$$\rho_b = \frac{\rho_b}{\rho_0}, \quad \bar{\rho}_u = \frac{\rho_u}{\rho_0}, \quad \bar{\lambda}_b = \lambda_b \frac{\rho^2}{k_{\text{off}}} \tag{45}$$

We further define a dimensionless reaction rate constant as ratio of binding and unbinding rates and an effective Peclet number

$$K \equiv \frac{k_{\text{on}}}{k_{\text{off}}}, \quad \mathcal{P}e \equiv \frac{L}{\tau_a}\frac{L}{D} = \frac{v_a L}{D}, \tag{46}$$

where $v_a \equiv L/\tau_a$ is a basal advection velocity by cortical flows.

For Rac and Rho surface densities, we choose $k_R$ and $k_\rho$ as characteristic concentrations and non-dimensionalize other parameters using previously defined time and lengthscales

$$\bar{R} = \frac{a}{k_\rho}, \quad \bar{\rho} = \frac{b}{k_R}, \quad \bar{\alpha}_{(0)} = \frac{\alpha_{(0)}}{k_\rho\, d_R}, \quad \bar{\beta}_{(0)} = \frac{\beta_{(0)}}{k_R\, d_\rho}, \quad \bar{d}_{R,\rho} = d_{R,\rho}\tau_a, \quad \bar{D}_{R,\rho} = D_{R,\rho}\frac{\tau_a}{L^2}. \tag{47}$$

The resulting dimensionless set of equations become

$$\frac{1}{\bar{d}_R}\frac{\partial \bar{R}}{\partial \bar{t}} - \bar{D}_R \bar{\nabla}^2 R = \Big(\bar{\alpha}_0 + \bar{\alpha}\,\text{th}(\bar{\rho}_b - \bar{\rho}_{th})\Big)\frac{1}{1+\bar{\rho}^2} - \bar{R} \tag{48a}$$

$$\frac{1}{\bar{d}_\rho}\frac{\partial \bar{\rho}}{\partial \bar{t}} - \bar{D}_\rho \bar{\nabla}^2 \rho = \Big(\bar{\beta}_0 + \bar{\beta}\,\text{th}(\bar{\sigma} - \bar{\sigma}_{th})\Big)\frac{1}{1+\bar{R}^2} - \bar{\rho} \tag{48b}$$

$$\bar{\sigma}_0 \nabla^2 \bar{x} - \frac{1}{\bar{\lambda}^2}\bar{\rho}_b\left(\dot{\bar{x}} - \bar{v}\right) = 0 \tag{48c}$$

$$\bar{\nabla}^2 \bar{v} + \bar{\nabla}\bar{\rho} - \frac{1}{\bar{\lambda}^2}\bar{\rho}_b\big(\bar{v} - \dot{\bar{x}}\big) = 0 \tag{48d}$$

$$\frac{1}{\bar{k}_{\text{off}}}\Big(\frac{\partial \bar{\rho}_b}{\partial \bar{t}} + \nabla(\bar{v}\cdot\bar{\rho}_b)\Big) = K\bar{\rho}_u - \bar{\rho}_b - \bar{\lambda}_b\,\bar{\rho}_b^3 \tag{48e}$$

$$\frac{1}{\bar{k}_{\text{off}}}\Big(\frac{\partial \bar{\rho}_u}{\partial \bar{t}} + \nabla(\dot{\bar{x}}\cdot\bar{\rho}_u) - \frac{1}{\mathcal{P}e}\nabla^2\bar{\rho}_u\Big) = -K\bar{\rho}_u + \bar{\rho}_b \tag{48f}$$

#### 5.4 Temporal persistence of polarization under optogenetic forcing

In this section, we investigate the temporal persistence of cell polarization when optogenetic forcing of Rac or Rho is applied, while keeping the basal production rates $\bar{\alpha}_0$ (for Rac) and $\bar{\beta}_0$ (for Rho) fixed.

In the main text (Fig. 6G), we identify three distinct regions in the ($\bar{\alpha}_0$, $\bar{\beta}_0$) parameter space:

1. A **biochemically bistable region**, where the system supports two stable steady states based on biochemical interactions alone.
2. A **mechanochemically polarizable region**, where persistent polarization arises due to feedback from mechanical processes, even though the biochemical system by itself is not bistable.
3. A **non-polarizable region**, where polarization cannot be sustained under any input.

We illustrate the influence of optogenetic Rho/Rac forcing across these three regions in Fig. 10, from left to right:

- **Non-polarizable region:** In this regime, no level of optogenetic input is sufficient to maintain a steady polarized state. This scenario is representative of neutrophil-like cells, which exhibit only transient polarization and ultimately return to an unpolarized state.
- **Mechanochemically polarizable region:** Starting from an unpolarized state, a sufficiently strong input—especially when both Rac and Rho are optogenetically activated—can drive the system toward persistent polarization. Notably, the response is asymmetric with respect to Rac and Rho inputs ($\bar{s}_R$, $\bar{s}_\rho$), reflecting differences in how Rac and Rho interact with the mechanical feedback. This asymmetry arises from differences in mechanical coupling strength, activation thresholds, and the nonlinearities in the biochemical-mechanical interaction. Importantly, the degree of asymmetry is sensitive to parameter changes within this coupling.

- **Biochemically bistable region:** Here, the system becomes ultrasensitive to small perturbations in Rac or Rho levels. Even minimal optogenetic stimulation can induce spontaneous polarization due to the inherent bistability of the biochemical network.

A key distinction between the mechanochemical model and a purely biochemical model (as depicted in Fig. 4) lies in their dependence on initial conditions. In the mechanochemical model, polarization occurs robustly regardless of the initial distribution of active GTPases. Once the system enters the bistable region, mechanical feedback ensures symmetry breaking and sustained polarization, whether the activated GTPase initially resides in a high or low concentration state.

In contrast, in a purely biochemical system, the outcome strongly depends on initial conditions. If optogenetic activation targets only the GTPase already in a high concentration regime, the system may not respond or evolve toward a new polarized state, underscoring the importance of mechanochemical feedback for robust symmetry breaking.

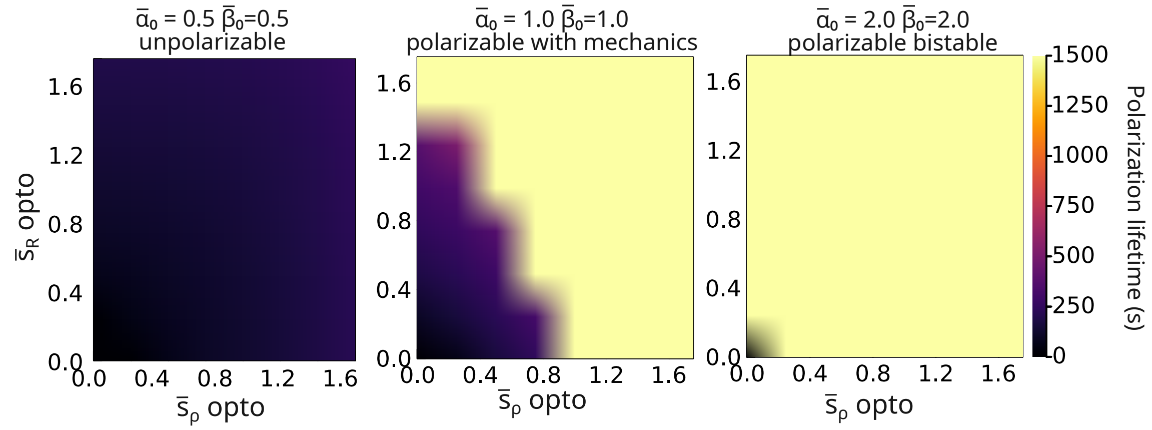


Figure 10: **Temporal persistence of cell polarization following optogenetic stimulation of Rac and Rho.** The heatmap shows how long the cell remains polarized after optogenetic exposure of a given intensity, with inputs $s_R$ (Rac) and $s_\rho$ (Rho). Simulations are based on the mechanochemical model without mass conservation of GTPases. Optogenetic stimulation is applied for 100 seconds at each end of the cell, after which the input is removed. The total simulation time is 1500 seconds; polarization durations of 1500 seconds or more are shown in yellow, as the colormap saturates at this upper limit.

### 6 Numerical Implementation

The equations are implemented using the finite-element method in a one-dimensional segment of size $L$, defined between the spatial coordinates $\xi = 0$ and $\xi = L$. The implementation is performed in the language *Julia* using the library *Gridap* [17, 18]. The code is available on a repository on Github.

We first present the weak formulation of the equations in one dimension and then the spatial and temporal discretization choices.

#### 6.1 Weak formulation

The weak form is derived from the strong form of the dimensionless equations (48) by multiplying each equation by an arbitrary test function $w$ and integrating over the spatial domain. For simplicity, we omit the overbars typically used to denote dimensionless variables.

$$\int_0^L w(\xi,t)\left[\sigma_0 \frac{\partial^2}{\partial \xi^2} x(\xi,t) - \frac{1}{\lambda^2}\rho_b(\xi,t)\big(\dot{x}(\xi,t) - v(\xi,t)\big)\right] d\xi = 0. \tag{49}$$

Integrating by parts we obtain

$$\begin{aligned}\int w\sigma_0 \frac{\partial}{\partial \xi}\frac{\partial x}{\partial \xi} d\xi &= \int k \tfrac{\partial}{\partial \xi}\Big(w \tfrac{\partial x}{\partial \xi}\Big)\partial \xi - \int \tfrac{1}{\lambda^2} \tfrac{\partial x}{\partial \xi} \cdot \tfrac{\partial w}{\partial \xi} d\xi \\ &= \sigma_0 \Big(w \tfrac{\partial x}{\partial \xi}\Big)\Big|_0^L - \int k \tfrac{\partial x}{\partial \xi} \cdot \tfrac{\partial w}{\partial \xi} d\xi \end{aligned} \tag{50}$$

The last line (50) corresponds the weak formulation of the problem.

Similarly for the cortex mechanics equation, we start from the integral

$$\int_0^L w\Big[\eta\frac{\partial^2 v}{\partial\xi^2} + \frac{\partial\sigma_a}{\partial\xi} - \chi\,\rho_b(\xi)\Big(v - \dot{x}\Big)\Big]d\xi = 0 \tag{51}$$

and will split the $\nabla^2$ by integration by parts

$$\int w\,\eta\nabla^2 v\,d\xi = -\int \eta\nabla w\nabla v\,d\xi + \Big(w\eta\nabla v\Big)\Big|_0^L. \tag{52}$$

Repeating the process for all the equations while taking adimensional equations, the system of weak equations is

$$\sigma_0\Big(w\nabla x\Big)\Big|_0^L - \int\Big[\sigma_0\nabla x\nabla w - w\,\frac{1}{\lambda^2}\,\rho_b\,\dot{x}\Big]d\xi = -\int w\,\frac{1}{\lambda^2}\,\rho_b v\,d\xi \tag{53a}$$

$$\Big(w\nabla v\Big)\Big|_0^L - \int\Big[\nabla w\nabla v - w\,\frac{1}{\lambda^2}\,\rho_b\,v\Big]d\xi = -\int\Big[w\,\nabla\rho + w\,\frac{1}{\lambda^2}\,\rho_b\,\dot{x}\Big]d\xi \tag{53b}$$

$$\int w\Big[\frac{1}{k_{\text{off}}}\Big(\frac{\partial\rho_b}{\partial t} + \nabla(v\cdot\rho_b)\Big) + \rho_b + \lambda_b\rho_b^3\Big]d\xi = \int w\,K\,\rho_u\,d\xi \tag{53c}$$

$$-\frac{1}{k_{\text{off}}\mathcal{P}e}\Big(w\nabla\rho_u\Big)\Big|_0^L + \int\Big(\frac{1}{k_{\text{off}}\mathcal{P}e}\nabla\rho_u\cdot\nabla w + w\Big[\frac{1}{k_{\text{off}}}\Big(\frac{\partial\rho_u}{\partial t} + \nabla(\dot{x}\cdot\rho_u)\Big) + K\,\rho_u\Big]\Big)d\xi = \int w\,\rho_b\,d\xi \tag{53d}$$

$$-D_R\Big(w\nabla R\Big)\Big|_0^L \int\Big(D_R\nabla R\cdot\nabla w + w\Big[\frac{1}{d_R}\frac{\partial R}{\partial t} + R\Big]\Big)d\xi = \int w\Big[\Big(\alpha_0 + \alpha\,\text{th}(\rho_b - \rho_{th})\Big)\frac{1}{1+\rho^2}\Big]d\xi, \tag{53e}$$

$$-D_\rho\Big(w\nabla\rho\Big)\Big|_0^L \int\Big(D_\rho\nabla\rho\cdot\nabla w + w\Big[\frac{1}{d_\rho}\frac{\partial\rho}{\partial t} + b\Big]\Big)d\xi = \int w\Big[\Big(\beta_0 + \beta\,\text{th}(\sigma - \sigma_{th})\Big)\frac{1}{1+R^2}\Big]d\xi. \tag{53f}$$

Here we separate the terms depending on the variable we will solve in the left hand-side from the other terms in the right hand-side.

#### 6.2 Spatial discretization

We use a cartesian discretisation and a scalar-valued Lagrangian finite elements space of order 1.

#### 6.3 Temporal discretization

The weak form of the partial differential equations (PDEs) above are solved in space at a given time. Time discretization for each variable $y$ is implemented as a simple forward Euler time-stepping

$$\frac{\partial y}{\partial t} \approx \frac{y_t - y_{t-1}}{\Delta t}, \tag{54}$$

where $\Delta t$ is the time-step.

We chose the time-step $\Delta t$ small enough to ensure proper convergence of the solutions of our equations. A range from $10^{-2}$s to $10^2$s was studied and we determined that 1s allows for good numerical accuracy while maintaining computational time acceptable.

#### 6.4 Parameters numerical values

There is a wide array of parameters to be set up in this model, which will be summarized here. The membrane tension $\sigma_0$ [N/m] can be measured from membrane tether pulling [19, 7]. The friction coefficient is estimated as $\mu = \chi\rho_0$, where $\chi \approx 10^{-6}$ Pa.s.m [20] is the drag coefficient for an individual linker within the membrane and $\rho = 10^{14}m^{-2}$ is a typical surface density of linkers [21].

| Notation | Quantity | Experimental Value | Ref(s). |
|---|---|---|---|
| $L = R\pi$ | Neutrophil cell half perimeter | 20$\mu$m | |
| $\sigma_0$ | Membrane tension | 10 pN/$\mu$m | [19] |
| $\eta_{3D}$ | Actomyosin viscosity | $10^4 - 10^5$Pa s | [10] |
| $\mu$ | Friction coefficient | 100 pN.s/$\mu m^3$ | [20, 21] |
| $T_0$ | Cortex thickness | 200nm | [22] |
| $\eta = \eta_{3D} T_0$ | 2D cortical viscosity | $10^5$ pN.s/$\mu$m | |
| $D$ | Diffusion of ezrin on the membrane | 0.003 $\mu$m$^2$/s | [23] |
| $k_{\text{on}}$ | Rate of ezrin binding | 5.0 1/s | [23] |
| $k_{\text{off}}$ | Rate of ezrin unbinding | 1.4 1/s | [23] |
| $\bar{\alpha}_0$ | Dimensionless rate of Rac activation | 0.1-10 | |
| $\beta_0$ | Dimensionless rate of Rho activation | 0.1-10 | |
| $d_R$ | Rate of Rac deactivation | 0.04 1/s | |
| $d_\rho$ | Rate of Rho deactivation | 0.04 1/s | |

**Table of values used for the different mechanical parameters.**
